# Self-avoidance dominates the selection of hippocampal replay

**DOI:** 10.1101/2024.07.18.604185

**Authors:** Caitlin S Mallory, John Widloski, David J Foster

**Author notes:** Corresponding authors contact: Caitlin Mallory,; David Foster.

## Abstract

Spontaneous neural activity sequences are generated by the brain in the absence of external input^1–12^, yet how they are produced remains unknown. During immobility, hippocampal replay sequences depict spatial paths related to the animal’s past experience or predicted future^13^. By recording from large ensembles of hippocampal place cells^14^ in combination with optogenetic manipulation of cortical input in freely behaving rats, we show here that the selection of hippocampal replay is governed by a novel self-avoidance principle. Following movement cessation, replay of the animal’s past path is strongly avoided, while replay of the future path predominates. Moreover, when the past and future paths overlap, early replays avoid both and depict entirely different trajectories. Further, replays avoid self-repetition, on a shorter timescale compared to the avoidance of previous behavioral trajectories. Eventually, several seconds into the stopping period, replay of the past trajectory dominates. This temporal organization contrasts with established and recent predictions^9,10,15,16^ but is well-recapitulated by a symmetry-breaking attractor model of sequence generation in which individual neurons adapt their firing rates over time^26–35^. However, while the model is sufficient to produce avoidance of recently traversed or reactivated paths, it requires an additional excitatory input into recently activated cells to produce the later window of past-dominance. We performed optogenetic perturbations to demonstrate that this input is provided by medial entorhinal cortex, revealing its role in maintaining a memory of past experience that biases hippocampal replay. Together, these data provide specific evidence for how hippocampal replays are generated.

## Main text

We recorded the activity of ensembles of place cells from the bilateral dorsal hippocampus of rats running laps on a linear track (*n=*11) or navigating within an open arena (*n=*6) (**Fig. 1a,f**). Rats traversed the linear track, pausing at each end to consume liquid food reward (mean±s.e.m. rewards per session: 53±3, *n=*52 sessions; **Fig. 1a-b**). In the open arena, rats were rewarded on alternating trials at a consistent location that was learnable across the session, or a random location that was unpredictable^17,18^ (‘Home trials’ and ‘Random trials’; mean±s.e.m. trials per session: 76±5, *n=*56 sessions; **Fig. 1f**). In some sessions (*n=*39) rats were additionally required to circumnavigate transparent ‘jail’ barriers^18^. Large numbers of co-recorded cells (mean±s.e.m. cells per session: linear track=82±05, open arena: 309±21) in conjunction with a memory-less, uniform-prior Bayesian decoding algorithm^19^ allowed us to accurately decode the rat’s position during movement (mean±s.e.m. positional decoding error per session, linear track: 2.9±0.1 cm, open arena: 3.4±0.2 cm). On the linear track, where place cells are directionally selective^20^, we additionally decoded the animals’ movement direction (**Fig. 1a**).

**Fig. 1.**
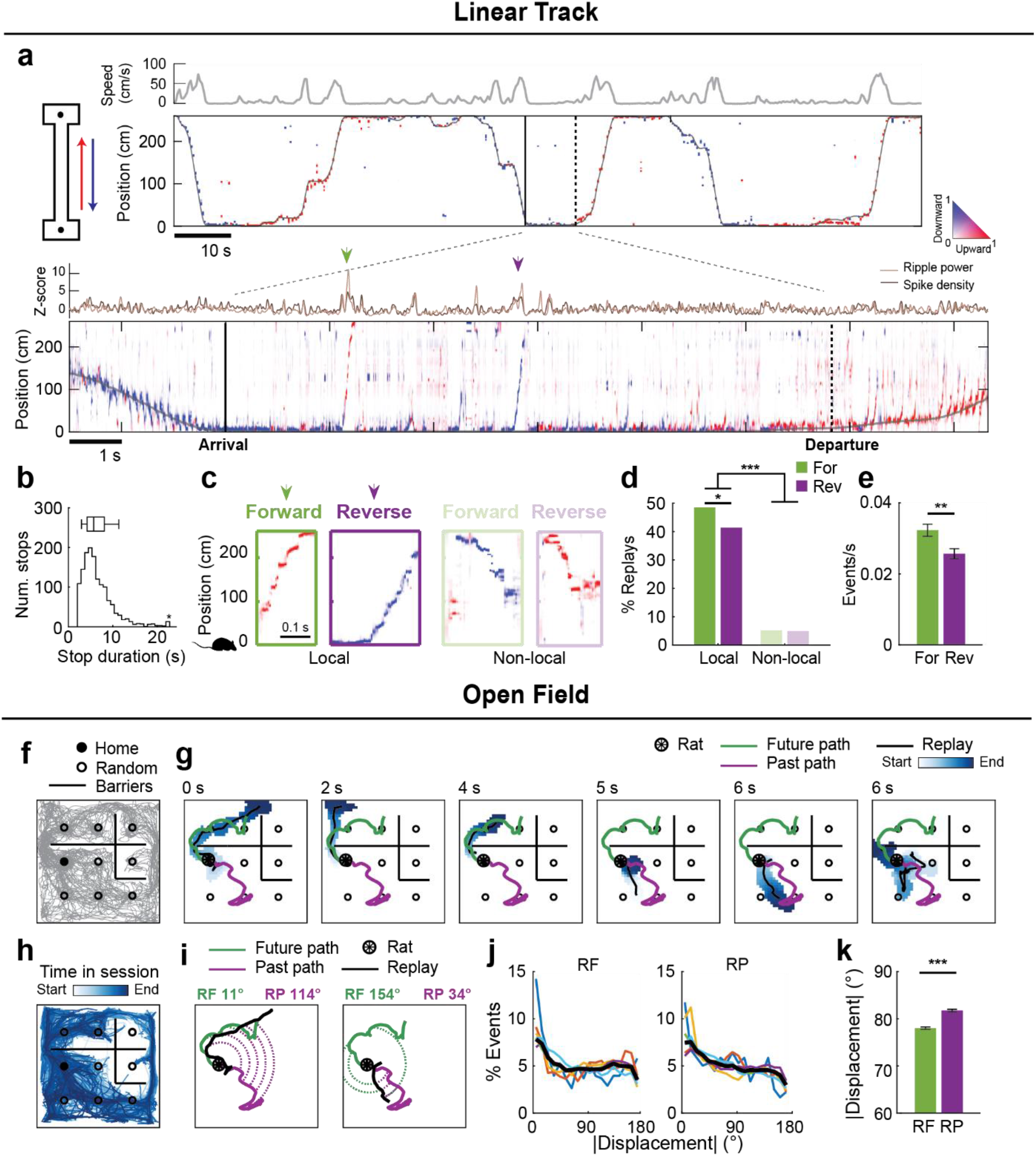
Linear track and open field replay events. **a**, *Left:* Schematic of the linear track. Circles: reward wells. *Right:* Posterior probability of position and running direction obtained from Bayesian decoding (400 ms bins) with the true position in gray. Color indicates the estimated running direction (% of time with direction correctly estimated, mean±SEM per session: 88.9±0.9%). *Inset:* posterior probability during one stopping period (20 ms bins). **b**, Stopping period durations. **c**, Example replays. **d**, Summary of replays occurring within the first 10 s of stopping. Local (*n=*849) versus non-local (*n=*96), *Z=*24.5, *P<*0.0001. Forward local (*n=*458) versus reverse local (*n=*391), *Z=*2.3, *P=*0.021; two-tailed z-tests. **e**, Forward and reverse replay rates per stopping period (mean±s.e.m., *n=*1642, *Z=*2.7, *P=*0.0058, Wilcoxon Signed Rank [WSR] test). **f**, Schematic of the open field arena. Behavioral trajectory (gray) and barrier configuration from one session is shown. **g**, Example replays from one stopping period. Time since stopping is shown at top. The posterior probability is color-coded to show elapsed time within the replay and was binarized to discard bins with low posterior (Methods). Black line: replay center-of-mass. Green: immediate future path of the rat. Purple: immediate past path (100 cm for each). **h**, The center-of-mass of all replay events in the session. **i**, Illustration of the calculation of angular displacement from the immediate future (RF) or past path (RP) for two example replays. Lower angles indicate greater replay:path similarity. See also Methods, and Fig. S4. **j**, Black: percent of replays as a function of angular displacement from future or past paths. Colored lines: individual rats. **k**, Mean±s.e.m. angular displacement from the immediate future or past path (*n=*36,677 replays, *Z=*-9.54, *P=*1.4-21, WRS test). \**P<*0.05, \*\**P*<0.01, \*\*\**P<*0.001.

Decoding on a fine timescale revealed abundant replay during reward-associated immobility (Methods, **Fig. 1a,g**). On the linear track, replays were classified as forward if the decoded heading direction aligned with the direction of replay motion across the track, and reverse if they were opposite (i.e., a trajectory depicting the animal running backwards)^9,10,19^ (**Fig. 1c**). We restricted analysis to replays representing motion away from the animal’s current location (‘local’, *n=*849), which made up the majority of events (**Fig. 1d**) and represented forward movement along the rat’s immediate future path or reverse movement along his immediate past path. Forward events occurred with slightly higher frequency than reverse events (**Fig. 1d-e**). In the open arena, we observed two-dimensional trajectories across the environment^17,18^ (**Fig. 1h**), many of which approximated the path taken immediately before or after stopping (**Fig. 1g**). To quantify the degree to which replays corresponded to adjacent behavior, we calculated the angular displacement between each replay event and the animal’s immediate past or future path, as previously^17^ (**Fig. 1i, Fig. S4**, Methods). Smaller angular displacement indicates greater similarity between the traversed and replayed paths. Consistent with previous findings, replays were overall more tightly aligned to an animal’s future path than his past^17,18^ (**Fig. 1j-k**).

### Forward/prospective replays precede reverse/retrospective replays

We observed a striking temporal order in the production of forward and reverse replays on the linear track (**Fig. 2**). Forward replays of the future path occurred shortly after stopping and almost always preceded reverse replays of the past (*n=*458 forward, 391 reverse replays; median [IQR] time since stopping, forward: 2.8 [1.7-4.6] s, reverse: 4.6 [3.4-6.4] s; *Z=*-10.8, *P*=5.2e-27, Wilcoxon Rank Sum [WRS] test). This forward-then-reverse ordering was evident across subjects, sessions, and individual stopping periods with multiple events (**Fig. 2a-b**). The incidence rates of forward and reverse replays varied across the stopping period and their difference revealed distinct windows marked by predominately forward (∼0-3 s, ‘forward window’) or reverse (∼3-10 s, ‘reverse window’) activity (**Fig. 2c-d)**. Theta power was strongly reduced during the forward window (**Fig. S1**), emphasizing that early forward replays are distinct from theta sequences^21–23^ seen during run. Moreover, we obtained similar results when requiring replays to coincide with a sharp-wave ripple (**Fig. S2**). Very few reverse replay events occurred in the forward replay window, prompting us to inspect the quality of the sparse reverse events detected shortly after stopping. The percentage of cells participating, spatial range, and precision of reverse replays were initially reduced compared to forward events, suggesting that timing constrains both the quantity and quality of replay content (**Fig. 2e**). We observed the temporal segregation of forward and reverse replays even in subjects with limited experience in an environment, indicating an intrinsically generated rather than learned organization (**Fig. 2f**). We did not observe consistent organization of replay content surrounding other behavioral transitions, emphasizing a unique significance of the run-to-rest junction (**Fig. S3**).

**Fig. 2.**
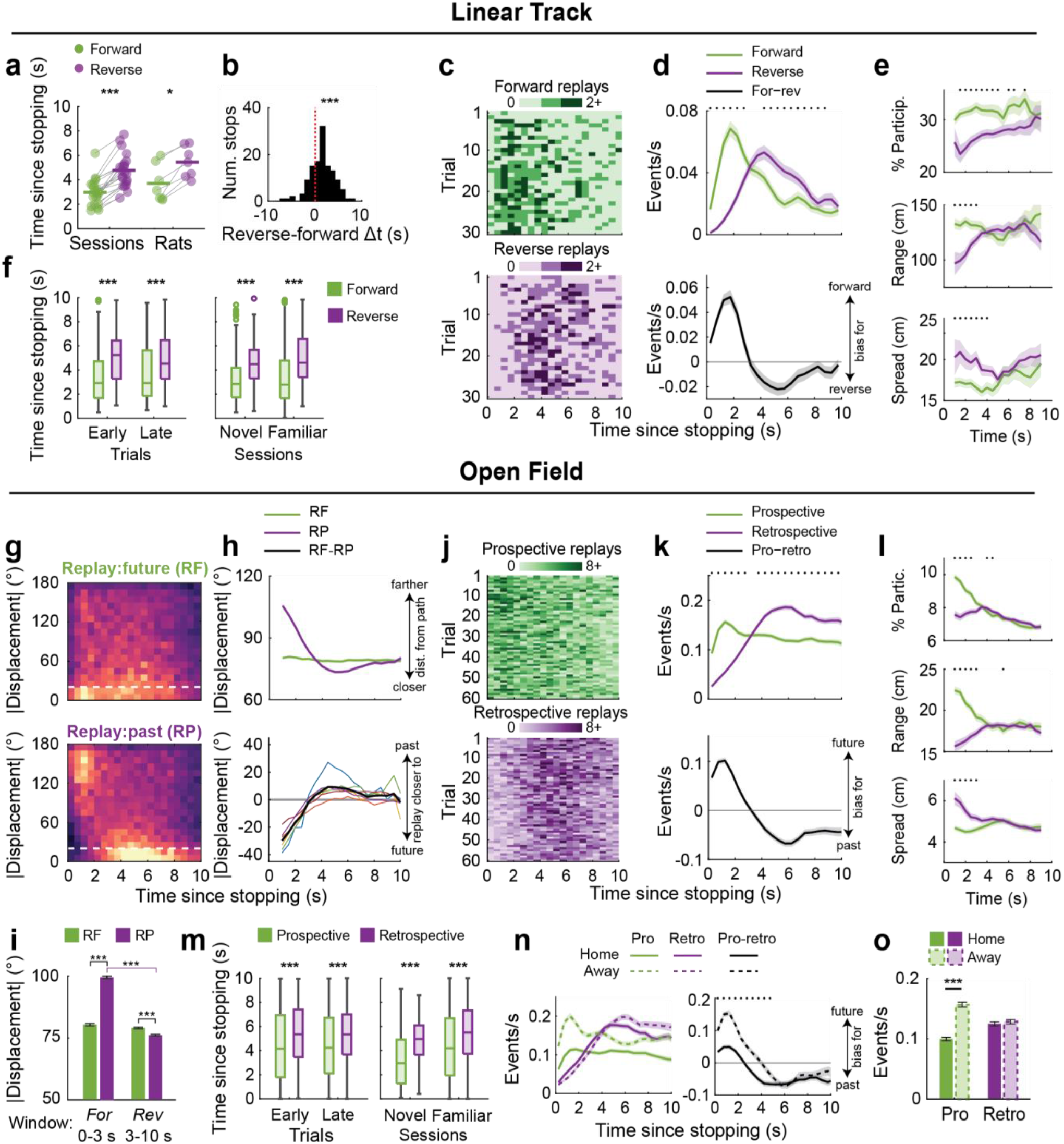
Forward/prospective replay precedes reverse/retrospective replay. **a**, Median time of forward or reverse replays (*n=*21 sessions/7 rats with at least 5 forward and 5 reverse replays detected; sessions: *P*=6.0e-5, rats: *P*=0.015, WSR). **b**, Time differences between the first forward and reverse replays within individual stopping periods (*n=*110, median [IQR] difference=1.4 [-0.2-3.0] s, *P*=2.5e-9, WSR). **c**, Replay counts, summed across sessions. **d**, Forward and reverse replay rates (*n=*a maximum of 1625 stopping periods; this number decreases over time due to variability in the time spent drinking). **e**, Replay properties (*n=*498 forward, 381 reverse replays in total, Methods). Lower values of spread indicate higher precision. **f**, Time of replays (early [trials 1-10]: *n=*46/61, *P*=4.4e-4; late [trials 30-40]: *n=*98/83, *P*=4.1e-5; Novel [first exposure]: *n=*105/106, *P*=9.4e-9; familiar: *n=*150/106, *P*=1.8e-19, WRS). Box plots: median, IQR, and range. **g**, Replay counts (max=200) at varying angular displacements from the future (RF, *top*) or past path (RP, *bottom*). **h**, Mean±s.e.m. angular displacement from the future or past path (*top*) and difference in angular displacement (*bottom*) (*n=*36,677 replays). Colors: individual rats. Black: rat average. **i**, Angular displacements within the forward (n=11,554 replays) or reverse windows (n=25123 replays) (interaction *P*=1.0e-4, shuffle; For window, RF v RP: *P*=4.5e-193; Rev window, RF v RP: *P*=1.2e-10; RF, For v Rev window: *P*=0.086; RP, For v Rev window: *P*=2.0e-315; WRS). **j**, Prospective (RF<20°) and retrospective replay (RP<20°) counts, summed across sessions. **k**, Prospective and retrospective replay rates (*n=*a maximum of 4,269 stopping periods). **l**, Replay properties over time (Methods; *n=*36,677 replays). **m**, Time of replays, as in (**f**) (early: *n=*637/705, *P*=2.6e-9; late: *n=*661/643, *P*=3.6e-11; novel: *n=*77/60, *P*=1.1e-4; familiar: *n=*4,752/4,951, *P*=4.3e-89; WRS). **n-o**, Prospective and retrospective replay rates on Home (*n=*2,147) and Away trials (*n=*2,122) over time (**n**) or averaged per stopping period (**o**). **o**, Interaction *P*=1.4e-4, shuffle; prospective rates, Home v Away: *P*=9.5e-22; retrospective rates, Home v Away: *P*=0.59, WRS. Except where otherwise stated, plots show mean±s.e.m. Asterisks in panels **d, e, h, k, l, n** indicate *P*<0.05, WSR tests. *P<*0.05*, *P<*0.001***

In the open arena, replay of the immediate future or past behavior unfolded across time in a highly consistent manner (**Fig. 2g**). Throughout the entire stopping period, many replays corresponded to the future path, consistent with an overall prospective bias^17,18^ (**Fig. 1k, Fig. 2g, top**). In contrast, replays were maximally distant from the past path just after stopping and became tightly aligned around 4 s (**Fig. 2g**, bottom). On average, replays aligned more tightly to the future path during the early forward window, and more tightly to the past path during the later reverse window (**Fig. 2h-i**). The temporal dynamics of this transition were highly consistent across individual rats (**Fig. 2h**, bottom, **Fig. S4**). We next restricted analysis to events that tightly aligned with behavior, classifying replays as prospective if they fell within 20° of the future path and retrospective if they fell within 20° of the past path. As on the linear track, the rate of prospective replays peaked shortly after stopping. The rate of retrospective replays was initially low but surpassed that of prospective replay after ∼3 s (**Fig. 2j-k**). Moreover, the quality of early retrospective replays was reduced (**Fig. 2l**), and the bias for prospective replays to precede retrospective replays was observable with little experience (**Fig. 2m**). While the early prospective bias at first appeared weaker in the open field, separate analysis of Home and Away trials revealed higher rates of prospective replay on the latter, which precede memory-guided navigation^17^ (**Fig. 2n-o**). The similarity in replay dynamics between linear track runs and open field Away trials likely reflects rats’ ability to anticipate their upcoming run in each case. Taken together, these data demonstrate a highly consistent temporal organization to replay content across behavioral tasks and environmental geometries.

### Replay avoids recently traversed or reactivated paths

We wondered whether the early, forward window in part reflects a tendency of replay to avoid the immediate past (**Fig. 3a**). Consequently, we analyzed the subset of open field trials (**Fig. 3b**) in which the rat retraced his steps, which resulted in overlapping past and future paths (past:future angular displacement <60°, Methods, **Fig. 3c**, top). If past avoidance were the dominant mechanism determining replay direction, we would expect early replays after stopping to avoid both the future and past paths (high angular displacement). However, if future preference were the dominant mechanism, early replays would be similar to both the future and past paths (**Fig. 3a**). Indeed, replays clearly avoided both the (overlapping) past and future paths for the first ∼3 s of stopping (**Fig. 3d-e** top). In contrast, when the past and future paths maximally diverged (past:future angular displacement >120°), early replays avoided the past and aligned with the future (**Fig. 3d-e**, bottom). We additionally quantified the tendency of replay to avoid the immediate past path to a greater degree than other potential paths intersecting the reward well at which the rat was positioned (**Fig. 3f**). Replays were more distant from the immediate past path than other experienced paths for seconds after stopping (**Fig. 3f**). Together, these data indicate the presence of a mechanism that prevents reactivation of the rat’s past path, even when that path is taken subsequently.

**Fig. 3.**
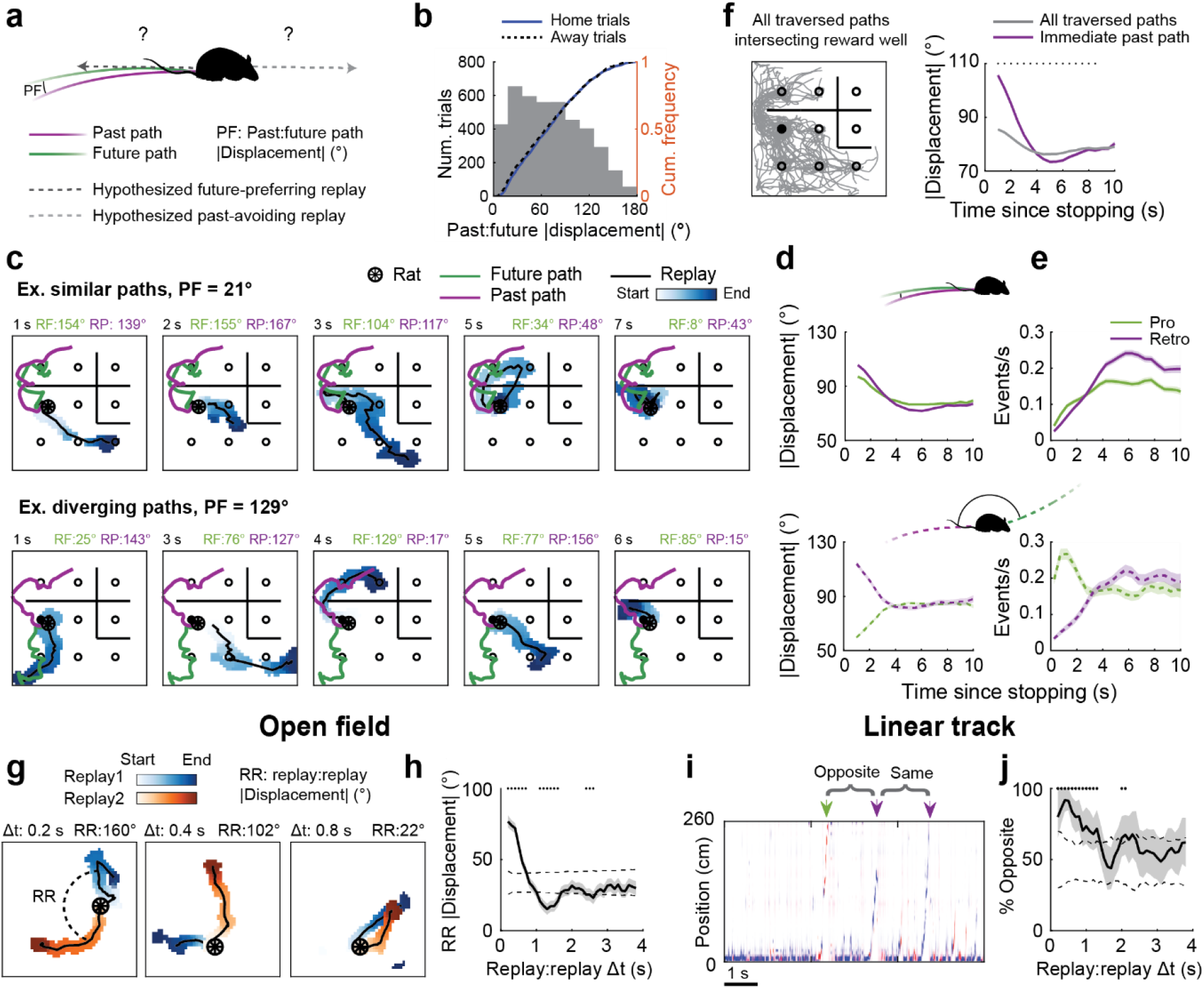
Replay avoids recently traversed or reactivated spatial paths. **a**, Illustration of the angular displacement between the past and future paths (PF), and hypothesized replays that could occur when this angle is small. **b**, Gray: PF angular displacements of all trials (*n=*4,269). PF did not differ between Home and Away trials (*P=*0.40, WRS). **c**, Replays from example trials with similar (PF<60°) or diverging (PF>120°) past and future paths. *Top left*: time since stopping. *Top right*: replay:future (RF) and replay:past (RP) angular displacements. **d**, *Top:* On trials with similar paths, both RF and RP decreased over time, demonstrating that replays early replays avoid both paths (RF: *r*(16,109)=-0.12, *P=*7.7e-57, RP: *r*(1,6109)=-0.19, *P=*3.1e-128; Pearson’s correlations). *Bottom:* on trials with diverging paths, RF increased over time while RP decreased (RF: *r*(5,369)=0.15, *P=*1.7e-28, RP: *r*(5,369)=-0.17, *P=*3.9e-35). **e**, Prospective and retrospective replay rates on trials with similar (*top*, max *n=*1880 trials) or diverging (*bottom*, max *n=*619 trials) paths. The rate of prospective replays during the 0-3 s forward window was reduced for trials with similar paths compared to those with diverging paths (*P=*2.5e-41, WRS test). **f**, *Left:* All traversed paths intersecting one (filled) well. *Right:* Angular displacement of replays relative to the immediate past path versus all other traversed paths (*n*=36,677 replays). Asterisks denote *P*<0.05, WSR tests. **g**, Example replay pairs. The time between and angular displacement between the two replays (RR) are shown at top. **h**, Angular displacement between two replays versus the time between them (*n*=42,345 replay pairs). Solid line, shading: mean±s.e.m. Dashed lines: 2.5% and 97.5% quantiles obtained by randomly permuting Δt. Asterisks indicate *P*<0.05, compared to shuffle. **i**, Example ‘opposite’ and ‘same’ replay pairs on the linear track. Green: forward. Purple: reverse. **j**, The proportion of ‘opposite’ replay pairs versus the time between them (*n*=389 replay pairs). Shaded region denotes the bootstrapped 95% confidence interval. Dashed lines: 2.5% and 97.5% quantiles obtained by randomly permuting Δt. Asterisks indicate *P*<0.05, compared to shuffle. **d-f**, plots show mean±s.e.m.

We hypothesized that replay would avoid a recently reactivated path in the same manner it avoids a physically explored path. We looked for evidence of this phenomenon in the open arena by asking whether two replays that occur in close temporal proximity are more likely to depict dissimilar spatial trajectories. To quantify the spatial similarity of two replay events, we calculated their angular displacement in the same manner as replay:path angular displacements (**Fig. 3g** Methods). Strikingly, replay:replay angular displacements were maximal when replays occurred in quick succession (**Fig. 3h**) and remained greater than chance for temporal distances up to ∼1 s (significance assessed relative to ‘shuffles’ in which the time between replay pairs was randomly permuted). Replays on the linear track showed a similar pattern of self-avoidance. We considered all events within a given stopping period and determined the percentage of replay pairs depicting opposing trajectories (i.e., forward followed by reverse, or reverse followed by forward, **Fig. 3i**). Replay pairs occurring within ∼1 s of one another were more likely than chance to depict opposing content (**Fig. 3j**). Together, these data reveal that replay direction is largely governed by avoidance of both recently traversed and reactivated spatial paths.

### MEC biases hippocampal replay toward retrospective sequences

Although this principle could potentially explain the early forward/past-avoidant window, it remained unclear how replay of past experience becomes predominant later into the stopping period. We reasoned that a memory of experience could be provided by medial entorhinal cortex (MEC), a primary source of cortical input to hippocampus^24^ that has been indirectly linked to reverse sequences^25^. We thus monitored hippocampal replay during optogenetic inactivation of MEC. To achieve widespread inhibition, we expressed the inhibitory opsin Jaws^26^ in bilateral MEC and delivered red light across the dorsal-ventral axis via tapered fiberoptics^27^ (**Fig. 4a-b, Fig. S5**). Two Jaws-expressing rats performed the open field navigation task. In experimental sessions (*n=*12), light was delivered to MEC during reward consumption (**Fig. 4c**). Control sessions (*n=*10) without light delivery were interleaved. While the overall rates of spike density events, sharp-wave ripples and replays were similar between MEC active or inactive trials (**Fig. S5**), MEC inactivation reduced the spatial range^28^ and cell recruitment of replays, particularly in the reverse window (**Fig. 4d**).

**Fig. 4.**
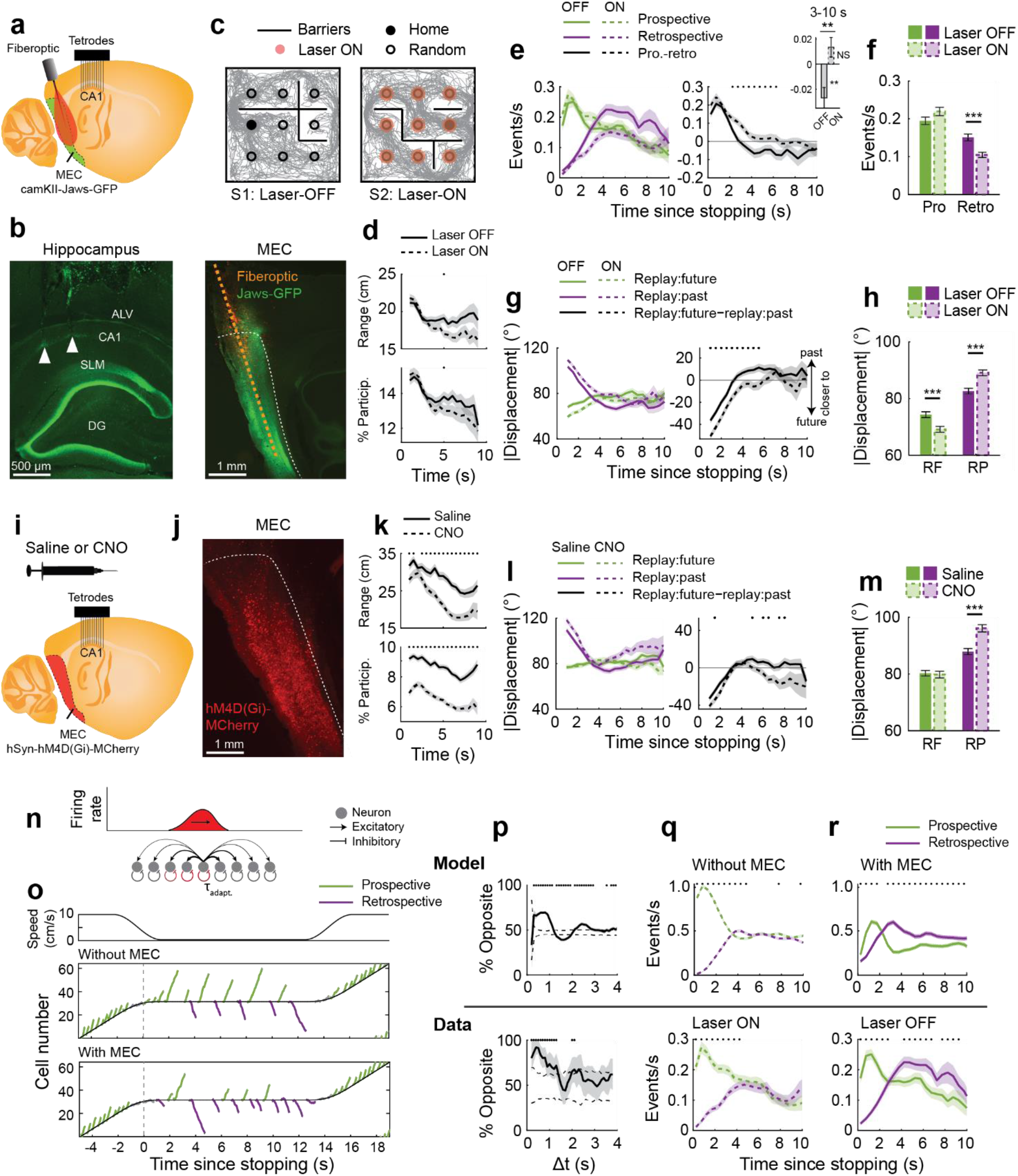
MEC activity produces a bias toward retrospective replay. **a**, Recording setup with optogenetic MEC inhibition. **b**, Histology. Arrows mark tetrode locations. DG: dentate gyrus; SLM: stratum lacunosum-moleculare; ALV: alveus of the hippocampus. **c**, Behavioral paradigm. **d**, Replay properties over time (*n*=2,778 OFF, 2,761 ON replays). OFF v ON within the reverse window (3-10 s), range: *P=*0.0068; % participation: *P=*0.0063 (WRS). **e**, Prospective and retrospective replay rates across time (*n=*maximum of 471 laser OFF, 484 laser ON trials). *Inset*: laser ON abolished the retrospective bias in the 3-10 s reverse window (prospective-retrospective rate differences, laser OFF: *P*=0.0030, laser ON: *P*=0.12, WSR; laser OFF v laser ON differences: *P*=0.0010, WRS). **f**, Prospective and retrospective replay rates within the first 10 seconds (interaction: *P=*1.0e-4, shuffle; OFF versus ON, prospective: *P=*0.20, retrospective: *P=*1.0e-4; WRS tests). **g**, Angular displacements (*n*=2,778 OFF, 2,761 ON replays) from the past or future path (*left*) and difference in angular displacement (*right*). **h**, Angular displacements within the first 10 seconds of stopping (interaction: *P=*1.0e-4, shuffle; OFF v ON, RF: *P=*2.4e-5, RP: *P=*4.0e-7, WRS tests). **i**, Recording setup. **j**, Expression of hM4D(Gi)-MCherry in MEC. **k**, Replay properties over time (*n=*2688 saline, 1922 CNO replays). Saline v CNO within the reverse window, range: *P=*3.4e-15; % participation: *P=*1.1e-42; WRS tests). **l**, Angular displacements from the past or future path (*left*) and difference in angular displacement (*right*). **m**, Angular displacements within the first 10 seconds of stopping (*n=*2688 saline, 1922 CNO replays (interaction, *P=*1.0e-4, shuffle; saline v CNO, RF: *P=*0.67, RP: *P=*8.8e-7; WRS tests). **e-m**, Plots show mean±s.e.m. **n**, The recurrent network model. **o**, Population activity during simulated track traversal. **p**, *Top:* The model produced self-avoiding (opposite) sequences at small time intervals. *Bottom:* Empirically observed self-avoidance (duplicated from **Fig. 3j**). Solid line, shading: mean±s.e.m. Dashed lines: 2.5% and 97.5% quantiles obtained by randomly permuting Δt. Asterisks indicate *P*<0.05, compared to shuffle. **q**, *Top*: Sequence rates produced by the adaptation-only model. *Bottom:* Empirically observed rates with MEC inactive (as in **e**). **r**, *Top:* Sequence rates produced by the modified model with simulated MEC (Methods). *Bottom:* Empirically observed rates with MEC active (as in **e**). **q-r**, plots show mean±s.e.m. \*\**P*<0.01, *** *P<*0.001.

We next asked whether MEC inhibition alters hippocampal replay content. Remarkably, MEC inactivation decreased the rate of retrospective replays and eliminated the retrospective bias typically seen in the later part of the stopping period (**Fig. 4e-f, Fig S5, Fig S6**). Examining the angular displacement of replays relative to the rat’s future or past path revealed similar effects: MEC inhibition increased the angular distance of replays from the past path, while decreasing the angular distance from the future path (**Fig. 4g-h**). We confirmed these results in an additional animal expressing the inhibitory DREADD hSyn-hM4D(Gi)-MCherry in bilateral MEC (**Fig. 4i-j**). Chronic inhibition via intraperitoneal injection of CNO reduced replays’ range, cellular recruitment, and alignment to the past path (**Fig. 4k-m**). We did not observe significant changes to task-performance with MEC inhibition (**Fig. S7**). MEC inhibition during intermittent periods of reward consumption on the linear track did not influence the frequency of replay events, but specifically decreased the quality of reverse replays (**Fig. S8**). Taken together, these results demonstrate that MEC activity promotes a bias for reverse/retrospective replay in the hippocampus.

### A network model recapitulates the temporal organization of replay

Finally, we explored possible mechanisms underlying these phenomena using a simple network model that has been applied to replay^29–32^. We simulated hippocampal place cell activity within a continuous attractor network, which supports a continuum of stable bump-like activity states^33–35^ (**Fig 4n**). Each neuron exhibits spike frequency adaptation, a form of slow timescale (∼1 s) feedback inhibition^36,37^, which destabilizes the activity bump, allowing it to sweep across the network in the absence of external input^29–32,38–40^. We first asked whether spike frequency adaptation could explain the tendency of replays to avoid self-repetition **(Fig 3g-j)**. We simulated exploration of a 1D track and initiated sequences with a brief, place-specific input at the animal’s current position (Methods, **Fig. 4o, Fig. S9**). During immobility, the model produced prospective and retrospective sequences that were self-avoiding over timescales consistent with our experimental findings (∼1 s, **Fig. 4p**). We next sought to identify whether the same mechanism could explain the more prolonged replay avoidance of the previously traversed path (**Fig. 3a-f**). At high running speeds, the model produced exclusively prospective sequences, consistent with theta sequences^22,23^ (**Fig. 4o**). Critically, as observed experimentally, replays continued to propagate forward for seconds after stopping, owing to the slow decay of the accumulated inhibition behind the animal. (**Fig. 4o-top, q**). Thus, within a recurrent hippocampal network, spike frequency adaptation at a fixed time scale can produce replay avoidance of recently reactivated or traversed paths across multiple timescales.

This adaptation-only model recapitulated the early window of past-avoidance, but not the later window of past-preference, shown experimentally to depend on MEC activity. We tested whether the window of past-preference could be recapitulated by including a facilitating input, presumed to arise from MEC, that accumulates in cells active during run and impinges on the circuit during immobility (Methods). As observed experimentally in the presence of MEC activity, retrospective replays became more frequent and indeed predominant once the experience-induced adaptation had dissipated (**Fig. 4o-bottom,r)**. Together, these results suggest a potential function of facilitating input from the cortex to bias the content of hippocampal replay.

## Discussion

Other models of replay generation are less compatible with the organization revealed here. Early models proposed that reverse replays could result from a slowly decaying activity trace following experience^9,10,41^. These models predict the highest rates of reverse replay immediately after stopping, in opposition to the observed window of past-avoidance. Other models based on recurrent neural networks with asymmetric synaptic weights do not easily generate reverse sequences and or predict replay self-avoidance^42,43^. The role played by spike frequency adaptation in our model, which has been observed in pyramidal cells of the rodent hippocampus^37^, could also be played by presynaptic depression^29^.

Similar models using spike frequency adaptation have been proposed to underlie hippocampal sequences observed during movement, including behavioral timescale ‘episode’ sequences^38,44^ and theta sequences^29,39,40^. These models successfully account for specific features of the latter, including backward-forward sweeps within individual theta cycles and alternating left-right sweeps across cycles^25,40,45,46^, raising the possibility that shared mechanisms operate across behavioral and sensory states.

In addition to the organization that results from spike frequency adaptation, we show for the first time that cortical input can exert top-down control over the content of awake hippocampal replay. Inhibition of MEC revealed its ability to impact the direction of hippocampal replay in addition to its length^28^. We modeled the MEC-driven enhancement of retrospective replay as an excitatory input current onto cells recently activated through experience. Such enhancement could result from calcium-mediated synaptic facilitation^47,48^ or persistent firing^49–52^ within spatially-selective MEC neurons^53–56^. Alternatively, this drive could be provided by retrospectively biased MEC replay^57,58^.

The organizing principles discovered here may underlie previously reported phenomena, including the enhanced replay of paths not-recently-taken^59–61^, or avoided entirely^62^. They also offer a potential explanation for the variability amongst prior works relating replay to ongoing behavior, highlighting the need to consider the time of events relative to stopping^13,63^. Interestingly, our findings are at odds with the predictions of a recent model of replay function, in which both forward and reverse replay drive the consolidation of reward predictions in downstream circuits^16^. Instead, the effective prioritization of forward replay may reflect its role in planning future behavior, for example, if planning an escape route is a priority over, or a precondition for, subsequent engagement in an episode of learning. Alternatively, postponement of reverse replay creates a temporal buffer between real and virtual experience, which could prevent interference between encoding and retrieval processes.

Refractory periods characterize neural circuits at multiple temporal and spatial scales^64–66^. Here we describe a novel refractory period in the production of internally generated neural sequences, with implications for the temporal structure of episodic memory retrieval during a memory-guided navigation task. This structure may promote exploration of alternative trajectories, as well as the avoidance of overtraining on highly valued options, and more generally the broadening of internally-generated training data in order to avoid bias and interference effects in network learning^67,68^. This may reflect a general principle governing internally-generated neural activity in multiple brain areas^1–8^.

## Methods

### Experimental model and subject details

All experimental procedures were in accordance with the University of California Berkeley Animal Care and Use Committee and US National Institutes of Health guidelines. Subjects were male Long-Evans rats (*Rattus norvegicus*; 3-9 months old, 450-550 g). Rats were housed in a humidity and temperature-controlled facility with a 12 h light-dark cycle. Before starting experiments, rats from the same breeding cohort were co-housed in pairs. Rats were single-housed at the start of experiments. All rats were implanted with microdrives containing dozens of tetrodes targeting the pyramidal layer of hippocampal region CA1 (see Drive design and surgery). Data presented here includes re-analysis of previously published data (Cohort 1: 5 rats performing linear track runs^69^; Cohort 2: 3 rats performing a goal directed navigation task^18^). New recordings of CA1 activity were obtained in 6 rats (Cohort 3). Of these, 4 rats were injected with an AAV encoding the inhibitory halorhodopsin Jaws^26^ bilaterally in medial entorhinal cortex (MEC) (‘Laser-Experimental’). 2 rats were injected with an AAV encoding the GFP bilaterally in MEC (‘Laser-Control’). One animal from each of the latter two groups was co-injected with an AAV encoding inhibitory DREADDS bilaterally in the MEC. In these animals, laser manipulations or administration of the DREADD agonist CNO were performed on separate days. Unless otherwise stated, analysis includes data from Cohorts 1 and 2 and control epochs (no-laser, no-CNO) from Cohort 3.

### Drive design and surgery

Virus injection and drive implant surgeries were performed separately, separated by 1–3 months to allow sufficient time for recovery and virus expression. Rats were anesthetized via 5% isoflurane inside an induction chamber before being transferred into a stereotaxic apparatus where they were maintained at 1-3% isoflurane. At the start of surgery rats received injections of buprenorphine (0.1 mg/kg), atropine (5mg/kg) and cefazolin (5mg/kg). In the virus injection surgeries, the bilateral MEC was injected with rAAV5/camKII-Jaws-KGC-GFP-ER2 (5.2×10^12 particles/ml), rAAV5/camKII-GFP-ER2 (5.2×10&12 particles/ml) or pAAV8/hSyn-HA-hM4D(Gi)-MCherry (2×10^13 particles/ml). Bilateral craniotomies (∼1 cm in diameter) were manually drilled above the lamdoid suture (∼10 mm posterior from bregma, 4.6 mm lateral of the midline) to expose the transverse sinus, used to identify the anterior-posterior position of MEC. A series of virus injections were made using a 10 μl World Precision Plus syringe controlled by a UMP3 micropump and Micro4 microsyringe pump controller. The needle was placed 0.2 mm anterior of the sinus, 4.6 mm lateral of midline, and angled 22° with the tip anterior. Injections of 250 nl each were made at the following depths relative to the brain surface: 5.2, 4.7, 4.2, 3.7, 3.2, 2.7, 2.2, 1.7 mm. Virus was delivered at a speed of 100 nl/min. Craniotomies were sealed with sterile duragel and dental sponge. In the subsequent drive-implant surgery, rats were implanted with microdrive arrays weighing 40-50 g and containing 40-64 independently adjustable platinum iridium (Neuralynx) tetrodes gold-plated to a final impedance of 150-300 MOhms. Drive cannulas were implanted bilaterally above the dorsal CA1 region of hippocampus (position from bregma: AP -4.1 mm, ML ± 2.7 mm) using previously described methods^17^. Virus-expressing rats were also implanted with tapered fiberoptics ^27^ (Optogenix Lambda Fibers, 0.39/200 [aperture, diameter]) in bilateral MEC. Fibers were angled 22° anterior and positioned at 0.3 ml anterior of the transverse sinus, 4.6 mm lateral of the midline. After lowering the fibers to a final depth of 5 mm relative to the brain surface, they were dental cemented in place. The fibers were custom-fabricated to emit light over the bottom 3 mm, thus delivering light to the majority of MEC, located ∼2-5 mm below the brain surface. Tetrodes were gradually lowered into the CA1 pyramidal layer over the course of 2-6 weeks, which was identified by the presence and polarity of sharp-wave ripples. The rats were allowed a week of recovery, after which food restriction and behavioral training resumed.

### Behavioral Tasks

Rats were habituated to daily handling and then food restricted to 85%-95% of their free-feeding weight. Rats were trained to run back and forth on a 1–2.6 m long, 6 cm wide, linear track to receive liquid chocolate milk reward (Nesquick). The tracks had two reward delivery wells at the ends. 6 rats additionally performed variants of the goal-directed, open field navigation task described in ^17^. Briefly, rats navigated on alternating trials to either a Home Well, whose location was held constant across the session, or a Random Well, whose location was randomly selected across trials. 2 rats performed the task as originally described^17^. Data includes 3 rats from a previous study^18^. The task requirements were similar to those of^17^, except that a) the rats were required to circumvent transparent jail barriers, whose locations in the arena changed from session to session and b) a green LED cued the location of the Random well, once filled. A detailed description of the apparatus and task protocol are available in^18^. 1 rat performed the task variation from^18^, but in the absence of LED cues.

### Manipulations of MEC Activity

A red laser (635 nm; Opto Engine LLC) was used in conjunction with tapered fiberoptics (0.39 NA Optogenix Lambda fibers, custom fabricated to emit light over the bottom ∼3 mm) to activate Jaws expressed along the dorsal-ventral axis of MEC. Optimal light emission via tapered fibers requires a light input with equal or greater numerical aperture^27^. We therefore delivered 0.57 NA light bilaterally to the tapered fiber optics as follows. The laser was connected via a 200 µm-0.22 NA patch cord to a custom rotary joint/splitter/NA converter (Doric Lenses). The splitter attached to two 200 µm-0.57 NA patch cords, which were attached to the ferrules of the tapered fiber optics (integrated in the Microdrive) at the start of each behavioral session. Laser power was calibrated to achieve a ∼20 mW at the fiber tip.

We first validated that this approach robustly and broadly inactivated MEC activity in anesthetized recordings. Rats (*n=*3) were anesthetized under isoflurane and placed in a stereotactic frame. A craniotomy was drilled above the MEC as described in *Drive design and surgery*. The tapered fiberoptic was glued parallel to a 64-channel silicon probe (Cambridge Neurotech L3 probes; channels spaced 50 µm apart, spanning a total distance of 3.15 mm) with ∼0.5 to 1 mm separating the fiberoptic and silicon probe. The fiber and probe were gradually lowered into MEC to a final depth of ∼5 mm over the course of 1 hour. Brief light pulses were applied (0.1 to 5 seconds each) for 5 minutes. Neural data was collected using the Digital Lynx data acquisition system (Neuralynx) and spike-sorted automatically with Kilosort2 (https://github.com/MouseLand/Kilosort^70^). Clusters were manually inspected and curated in Phy (https://github.com/cortex-lab/phy). For each cell we computed a laser modulation index as (FR_OFF_ - FR_ON_)/(FR_OFF_ + FR_ON_) where FR_OFF_ and FR_ON_ are the cell’s average firing rate during laser-OFF and laser-ON epochs. This index ranges from -1 (suppression) to 1 (excitation).

During behavioral experiments, continuous light was delivered during reward consumption. The laser was TTL-activated and controlled manually via a keystroke. For manipulations of MEC activity on the linear track, the laser was turned on on alternating laps either during reward consumption-only (*n=*27 sessions), or reward consumption and running (*n=*7 sessions). Thus, half the reward-epochs within a session were accompanied by light. For manipulations of MEC activity in the open field, behavioral sessions were conducted with either laser-OFF (*n=*11 session) or laser-ON (*n=*12 sessions). During laser-ON sessions, the laser was on during all periods of reward consumption at either the Home well or Random Wells, and off during movement.

The effects of laser-mediated MEC inactivation on hippocampal replay were further confirmed via DREADD-mediated inhibition of MEC in one rat. The subject performed 19 sessions of the goal-directed navigational task (up to two 2 sessions daily, separated by at least 4 hours). 20 minutes prior to each recording session, the rat received an intraperitoneal injection of either the DREADDS agonist clozapine N-oxide (CNO; 3mg/ml; 10 sessions) or an equivalent volume of sterile saline (9 sessions). On days in which the subject performed two sessions, CNO was administered in the second session.

### Behavioral analysis

Rat position was tracked by an overhead camera and sampled at 30 Hz. Position data was determined from red and green LEDS mounted on the headstage. Position and velocity data were smoothed using a Butterworth filter (second order with a cutoff frequency of 0.1 samples/s using MATLAB’s butter function). On both linear track and open field sessions we aligned replays to movement cessation/the start of reward consumption (Fig. 2–4) or end of reward consumption (Fig. S3). For each trial, movement cessation/the start of reward consumption was taken as the first time the rat entered within 5 cm of the filled reward well at a speed of 3 cm/s or less. The end of the reward consumption was taken as the first time the rat exited the reward well at a speed of 5 cm/s or greater. On linear track sessions we additionally aligned replays to run onset– the time at which the rat departed from the reward platform (Fig. S3). This moment can differ from that marking the end of reward consumption because following reward rats occasionally hesitated before running across the track again. Run onset was taken as the moment the rat exited the reward platform area (30 cm) at a speed of 10 cm/s or greater.

#### Determination of past and future paths

The past and future behavioral path of the rat was computed for each stopping period (reward consumption at either the Home or Random wells). We defined the rat’s past path as the 180 cm of path preceding drink onset, and the future path as the 180 cm of path proceeding departure from the reward zone. Thus, all replay events within a stopping period were compared against the same 2 behavioral paths.

#### Angular displacement between past and future paths

For each stopping period in the open field, the angular displacement between the rat’s past and future paths was computed in a manner similar to replay:path angular displacement (see below). A series of centric circles was centered on the position of the rat at the start of the stopping period, and the minor arcs formed by the intersections of these rings and the two paths (past and future) were averaged. Thus, for each stopping period we obtained a single angle describing the similarity between the past and future path.

### Data Acquisition

Neural data was collected using Neuralynx/Cheetah or Spike Gadgets/Trodes acquisition systems (sampling rate 32.5 kHz, 30 kHz respectively). Spikes exceeding 50 µV were clustered manually (xclust2, M.A. Wilson, MClust, David Redish) or automatically (Mountainsort^71^). Automatically-determined clusters were accepted if they passed visual inspection and met the following criteria: noise overlap < 0.03, isolation > 0.5, peak SNR > 1.5^71^. LFP was digitally filtered between 0.1 and 500 Hz and recorded at 3,255 Hz (Neuralynx) or 1500 Hz (Spike Gadgets).

### Spike density and sharp-wave ripple amplitude

Population spike density was computed by totaling the number of spikes from all cells in 1 ms non-overlapping time bins. The LFP from a selected tetrode in the pyramidal layer with visually identified sharp-wave ripples was band-pass filtered between 150 and 250 Hz, and the amplitude was computed as the envelope of the Hilbert transform. Spike density and ripple amplitude were both smoothed with a Gaussian kernel (12.5 ms standard deviation [SD]). Periods in which the rat’s speed was below 5 cm/s were z-scored. Peaks in the z-scored spike density or ripple amplitude traces exceeding 3 SD above the mean were identified as spike density or ripple events. The start and end time of each event was defined as the times on either side of the peak at which the z-scored signal crossed the mean.

### LFP spectral analysis

To visualize how the LFP power in different frequency bands varies across a trial, we constructed time-frequency spectrograms using Morlet wavelet convolution. For each trial (−2–10 seconds, aligned to reward consumption), the LFP signal from a single tetrode located in CA1 was convolved a series of 200 Morlet wavelets (1-251 Hz, logarithmically spaced; the number of cycles varied from 4 to 10 for an optimal balance of temporal and frequency precision^72^ to obtain a time-varying estimate of power. The power at each frequency was separately z-scored over time to facilitate comparison across sessions and animals. An average time-frequency spectrogram was computed for each behavioral session by averaging across trials.

### Place fields

The rat’s position and speed at each spike time were computed through linear interpolation (*interp1* in MATLAB). Place fields were calculated from periods of movement (speed exceeding 5 cm/s). Linear track passes were separated by running direction to construct unidirectional rate maps. Positions were binned into 2 cm bins (linear track) or 2×2 cm square bins (open field). Unsmoothed rate maps were computed as the number of spikes per bin normalized by occupancy. On the linear track, raw maps were smoothed with a gaussian kernel (SD 2 cm). In the open field, unvisited bins were set to zero and the raw maps were convolved with a 2D isotropic Gaussian kernel (8 cm SD)^18^.

### Bayesian decoding of position

The posterior probability (*P*) of the animal’s position (pos) is given by Bayes rule (assuming Poison firing statistics, independence between neurons, and a uniform prior over position^19^.

The posterior probability of the animal’s position (pos) across *L* total position bins given a time window (*τ*) containing neural spiking (spikes) is

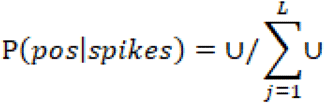

where on the linear track

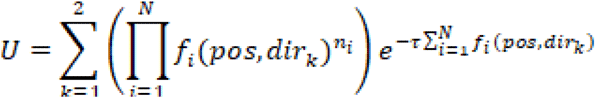

*f*_*i*_(*pos,dir*_*k*_) is the *i*-th place field in a running direction, and *n*_i_ is the number of spikes emitted by the *i*^*th*^ cell in the time window (*τ*).

In the open field,

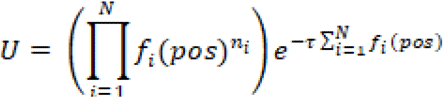

Where *f*_*i*_ (pos) is the position tuning curve of the *i*^*th*^ unit, and *n*_i_ is the number of spikes emitted by the *i*^*th*^ cell in the time window (). The posterior probability was computed for all bin locations x_j_ where 1≤j≤L and L is the total number of spatial bins.

During movement, position was estimated from non-overlapping 400 ms windows and compared to the true position to determine the decoding error. Sessions for which the average decoded positional error exceeded 10 cm were discarded from further analysis. During immobility, position was estimated in smaller temporal windows (linear track: 20 ms overlapping by 5 ms^73^, open field: 80 ms overlapping by 5 ms^18^).

### Replay detection

#### Linear Track

The posterior probability over position and movement direction was computed for each candidate replay event (spike density events). Replay events were required to satisfy the following criteria: >66% of the posterior probability located in one of the directional maps, weighted correlation >0.6, max. jump distance<40% of the track, spatial coverage > 20% of the track and at least 10 cells participating ^69,73^. We verified our results using SWRs as candidates (Fig. S2).

*Weighted correlation*: decoded probabilities (*P*) were assigned as weights of position estimates to calculate the correlation coefficient between time (*T*) and decoded position (*Pos*):

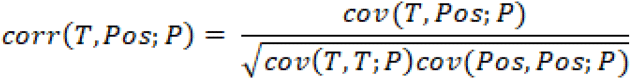

where weighted covariance between time and decoded position is

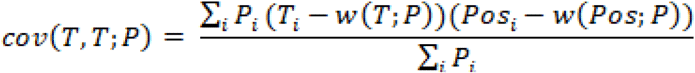

and weighted means of time and decoded position are

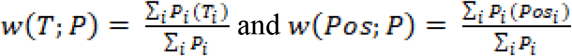

*Maximum jump distance* was defined as the maximum distance between peak decoded positions in adjacent time windows for each candidate event normalized by the track length. *Posterior spread*: as a measure of posterior spread, for each replay we computed the average 95% confidence interval across all time bins. Within a time bin, the 95% highest posterior density was computed as the smallest continuous spatial interval containing 95% of the posterior probability, normalized by the track length. The *spatial coverage* (*range*) of a replay was calculated as the distance between the minimum and maximum positions decoded within the event.

#### Open Field

To identify replay, we decoded the posterior probability over position over the entire session using 80 ms time windows overlapping by 5 ms^18^. Replay events were defined as stationary epochs in which the posterior was sharply tuned (small posterior spread) and moved smoothly (small *COM jump size* across time steps)^18^. A subsequence was defined as a set of contiguous bins meeting the following criteria: rat speed <5 cm/s, posterior spread (*m*<0.0048*L cm*) and posterior COM jump size 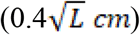 where *L* is the total number of spatial bins. Neighboring subsequences were merged if they were separated by less than 20cm and 50ms. The final sequence was required to have a duration greater than 50ms.

#### Posterior center-of-mass (COM)

Define *P*_*j*_ = *P*(*x*_*j*_|spikes) and let x_j_ =(x_j_,y_j_) be the coordinates of the *j*^*th*^ spatial bin. The coordinates of the *posterior center-of-mass* (COM) were given by:

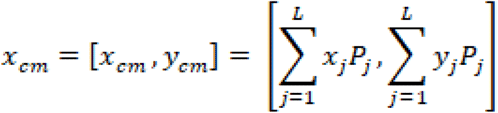

where *L* is the total number of spatial bins.

*Posterior spread* was defined as the square root of the second moment of the posterior:

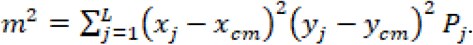

*Posterior COM jump size* was defined as the L2 norm of the difference vector between consecutive posterior center-of-mass estimates, 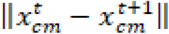.

*Spatial range* of a replay event was defined as the L2 norm of the difference vector between first and last posterior center-of-mass estimates, 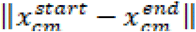.

Except in Figure S2, we did not require events to overlap with SWRs or SDEs or impose a minimum spatial coverage threshold^17,18^. These relatively permissive criteria were selected to avoid detection heuristics that may inadvertently bias the content of replay discovered^74,75^.

### Analysis of replay statistics

#### Replay self-avoidance

Fig. 3 reports that replays exhibit a tendency to avoid repetition over short time intervals. On the linear track, we examined the likelihood that two linear track replays depict ‘opposite’ content as a function of the elapsed time between them (Δt). ‘Opposite’ pairs consisted of one forward and one reverse replay. ‘Same’ pairs consisted of either two forward or two reverse replays. We considered all pairs of locally initiated replays occurring within 4 seconds of one another in the same stopping period. Replay pairs were binned according to their Δt (400 ms bins shifting by 100 ms). For each time bin we computed the proportion of Opposite replay pairs. 95% confidence intervals on these proportions were computed by bootstrapping. To determine which time bins contained Opposite proportions that differ significantly from chance, we compared the observed proportions to those obtained by randomly permuting Δt. For each shuffle iteration (*n=*5000 shuffles) we computed the proportion of Opposite-pairs in each time bin. Significant time bins are those for which the observed proportion falls outside the inner 95% quantile of the corresponding shuffled proportions.

In the open field, we compared the angular displacement between two replay events within the same stopping period as follows. A series of concentric circles was centered on the position of the rat at the start of the first replay event, and the minor arcs formed by the intersections of these rings and the two replay trajectories were averaged. Thus, for each replay pair we obtained a single angle describing the similarity between the two replay trajectories. Only locally initiated replays (starting within 16 cm of the rat’s physical position) were included in this analysis. Fig. 3 shows the mean angular displacement between two replays as a function of the time elapsed between them (Δt). To construct this plot, replay-pairs were binned according to their (Δt) (400 ms time bins, shifting by 100 ms) and the mean angular displacement was calculated. The significance of each time bin was determined by comparison to a shuffle in which Δt was randomly permuted across replay pairs. For each shuffle iteration, the Δt for all replay pairs was randomly permuted and a Δt versus angular displacement plot was regenerated. This procedure was repeated 5000 times to obtain a shuffled distribution of angular displacements over time. Time bins were significant if the observed mean angular displacement was outside the inner 95% quantile of the corresponding shuffled data.

#### Replay displacement from past or future path

For open field replays, the angular displacement from the rat’s past or future path was computed in a similar manner as previous work^17^. For each replay event, a series of concentric circles with increasing radii (14 to 180 cm, increasing in 2 cm increments) were centered on the rat’s position at the time of the replay. For each circle we found the minor arc formed by its intersection with the replay and relevant behavioral path (future or past; computed for each stopping period). The arcs from all circles were averaged. Thus, for each replay event we obtained 2 values: the mean angular displacement relative to the past path, and the mean angular displacement relative to the future path. For cases in which there were multiple intersections of the circle and either the replay or path, the intersection that occurred closest in time to the replay was used. In Figures S4 and S6, we additionally considered the angles formed by each crossing point, as in^17^, rather than averaging the angles per replay.

#### Classification as prospective and retrospective

Open field replays were classified as prospective and retrospective if their angular displacement from the future or past path was <20°, respectively. The categories were mutually exclusive (i.e., events within <20° of both the past and future paths were excluded), except in Fig. 3e, where we investigated the content of replay events on trials in which the past and future paths were minimally separated. In that case, mutually exclusive categories would exclude many of the replays that occurred when the past and future paths were separated by <20°. Thus in that case, a replay was considered prospective if it was <20° of the future and closer to the future path than the past path, and retrospective if it was <20° of the past path, and closer to the past path than the future path.

#### Calculation of replay rates over time since stopping

We computed replay event rates over time as follows. For all rewarded stops, the number of replay events within nonoverlapping 0.5 s windows was calculated and converted into rate by dividing the replay count by the time interval (0.5 s). Rates were calculated for the time interval [0 10 s], where 0 is defined as the moment the rat first entered within 5 cm of the reward at a speed of < 3 cm/s. If the rat did not remain stationary for the full 10 s, time bins in which the animal was no longer stationary at the reward zone were set to NaN. For each trial, the raw replay rate across time was smoothed by convolving with a Gaussian kernel (SD 0.5 s). Figures report the mean±s.e.m. across trials. Spike density event rate and ripple rate were computed similarly.

#### Calculation of replay properties over time since stopping

Replay metrics, including angular displacement from the past or future path, were calculated at different times since stopping (2 s bins shifting by 0.5 s). The mean±s.e.m. is shown for visualization. Statistical analysis was performed at each time bin using WRS tests.

*Away-events* were defined as replays that occurred while the rat drank at a Random well within a Random trial. *Home-events* were defined as replays that occurred while the rat drank at a Home well within a Home trial.

## Simulations of replay sequences

Hippocampal sequences were simulated using a single-bump continuous attractor network (CAN) where the movement of the bump was driven by spike-frequency adaptation^32,38,76,77^.

Given a summed input current to the *i*^*th*^ neuron, the instantaneous firing rate of the cell was *f*(*I*_*i*_(*t*)), with the neural transfer function given by 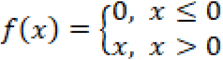.

Based on this time-varying input, neurons fired spikes according to a Poisson point process with a coefficient of variance of 1.

The input rate for the *i*^*th*^ neuron was a combination of internal and externally-derived inputs, each modulated at theta frequency:

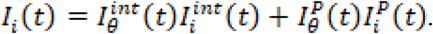

The “internal” input was given by

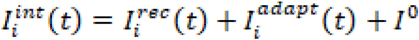

where 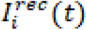 is the recurrent input derived from other cells (see below), 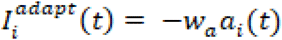 the adaptive inhibitory input (*w*_*a*_ is the strength of depression) which models the effects of slow calcium-dependent potassium currents^36^, and *I*^*0*^ is a small positive constant bias common to all cells. The recurrent input was

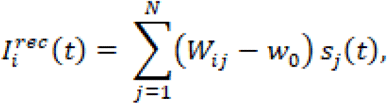

where *w*_*ij*_ are the excitatory recurrent weights, *w*_0_ is the strength of recurrent inhibitory feedback, and *N* is the number of neurons. To specify the recurrent weights, neurons were organized into a 1D periodic array in the neural sheet, where the location of the *i*^*th*^ neuron was given by *x*_*i*_ Let *w*_*ij*_ be a set of translation-invariant symmetric weights with Gaussian shape that depend on the distance between cells in the neural sheet:

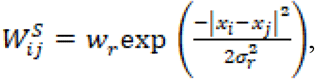

where *w*_*r*_ and *σ*_*r*_ control the strength and spatial extent of the connectivity, respectively. The synaptic activation dynamics for the *i*^*th*^ neuron,*s*_*i*_(*t*), was given by

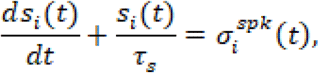

where τ_*s*_is the synaptic time constant and

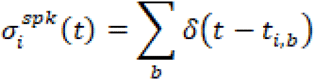

is the cell’s spike train (*t*_*i,b*_ specifies the time of the *b*^*th*^ spike of the neuron and the sum is over all spikes of the neuron). The adaptation dynamics for the *i*^*th*^ neuron, *a*_*i*_(*t*), was given by

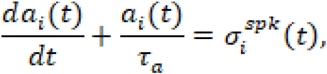

where τ_*a*_ is the time scale of adaptation.

To generate the corrective place inputs, a rat trajectory was simulated, starting at the origin, by generating a speed profile that was a sinusoid with period *T*_*transition*_ seconds but modified so that the rat dwelled in the rest state (zero velocity) for *T*_*rest*_ seconds and in the maximum velocity state for *T*_*run*_ seconds (Fig. S9). This speed profile was then integrated to generate a spatial trajectory. The place input into the *i*^*th*^ cell was given by a 1D Gaussian centered on the rat’s location at time *t* and width σ_*rat*_ and amplitude *w*_*rat*_:

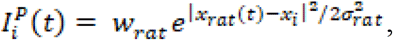

where *x*_*rat*_(t) is the rat’s position at time *t*.

Lastly, to mimic the transient suppression of sequences during the transition from running to stopping that is seen in real data, we defined transitions to and from running as the timepoints when the rat’s speed rose above or dipped below 3 cm/s. Next, we convolved an inverted Gaussian with a 1.5 s standard deviation around each transition point, then renormalized the curve such that it ranged between [0,1]. This curve was then used to modulate the strength of the adaptation, *w*_*a*_, over time.

### Theta modulation

Both the internal and place inputs were modulated at theta-frequency by a rectified sinusoid with a frequency that was a function of rat speed

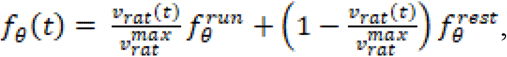

where 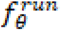 and 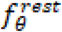 represent the maximum and minimum in the frequency of sequence generation and *v*_*rat*_(t) and 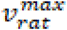 are the rat’s instantaneous and maximum running speeds, respectively (see below) (Fig. S9). To introduce variability in theta cycle lengths, theta periods were sampled iteratively from the distribution 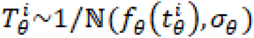 where ℕ is a normal distribution and 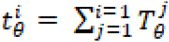 is the cumulative sum of all preceding theta periods. The theta modulation 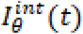 was then defined as follows: For the *i*^*th*^ theta interval of length 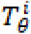 at time 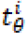,

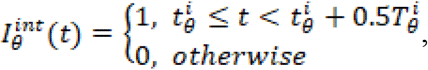

which amounts to a rectified sinusoid with a duty cycle of 50% of the theta period. Likewise, the place inputs were also modulated at theta-frequency:

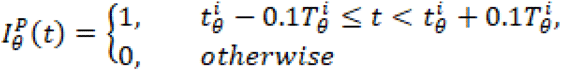

which amounts to a rectified sinusoid with a duty cycle of 20% of the theta period but also shifted in time in order to allow the place inputs to occur briefly around the time of the release of the internally-driven dynamics of the network (Fig. S9).

### MEC-driven facilitation

A third MEC-driven facilitating input was added to each neuron’s total input

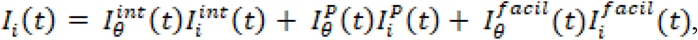

where 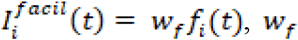, is the strength of facilitation and the facilitation dynamics *f*_*i*_(*t*) was governed by

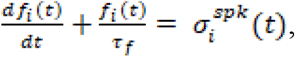

with σ_*f*_ being the time scale of facilitation. The theta modulation 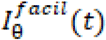 is assumed to have identical dynamics as 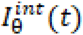 During running periods (*v*_*rat*_ > 10 cm/s) facilitating input was turned off (*w*_*f*_ = 0), whereas during stopping periods (*v*_*rat*_ ≤ 10 cm/s) the facilitation dynamics no longer received input from spiking:

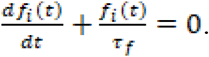

## Simulation parameters

Integration was by the Euler method with step size equal to 0.5 msec. Other network parameters were:

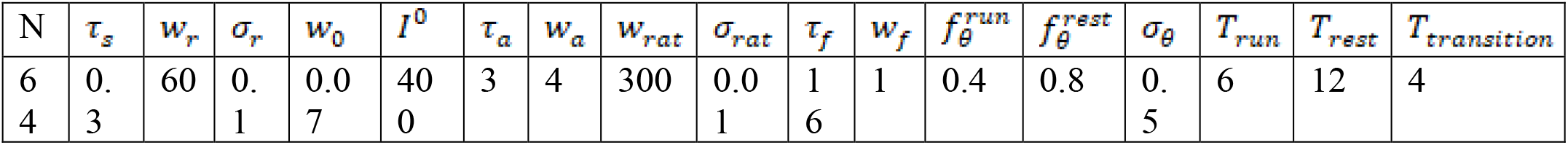

Temporal parameters are in units of seconds; amplitudes (*w*’s) are dimensionless; *I*^0^ is in units of spikes/sec.

## Statistical analysis

All statistical tests were two-tailed. In cases where data were not normally distributed, non-parametric statistical tests were used to estimate significant differences between groups (Wilcoxon rank sum or Wilcoxon signed rank tests). To test for an interaction between two factors when data were not normally distributed, a shuffling procedure was performed. The data fell into 4 groups: (a_1_, a_2_, b_1_, b_2_). We compared the observed difference of differences, 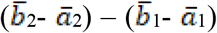, to a shuffled distribution (n=10000 shuffles) obtained by permuting the subgroup labels (for example, a_1_ and a_2_).

## Data availability

The datasets generated during the current study will be available prior to publication.

## Code availability

Code for reproducing the analyses in this article will be made available prior to publication.

Methods references

## Acknowledgements

The authors thank Sandro Romani for helpful discussions, Matthew Kleinman, Mark Plitt and Thomas Elston for comments on the manuscript, and Charlie Walters for assistance in designing the Microdrive. This work was supported by the National Institute of Health grants NS113557 and MH103325 (D.J.F.) and a Howard Hughes Medical Institute Hanna H. Gray Fellowship (C.M.).

## Author Contributions

C.M. and D.J.F. conceived the design of the study. C.M. performed the experiments and analyzed the data. J.W. contributed previously published data and performed the modeling work. C.M. and D.J.F. wrote the manuscript.

## Supplementary Information

is available for this paper.

## Competing interests statement

Authors declare that they have no competing interests.

## Supplemental Figures

**Fig. S1.**
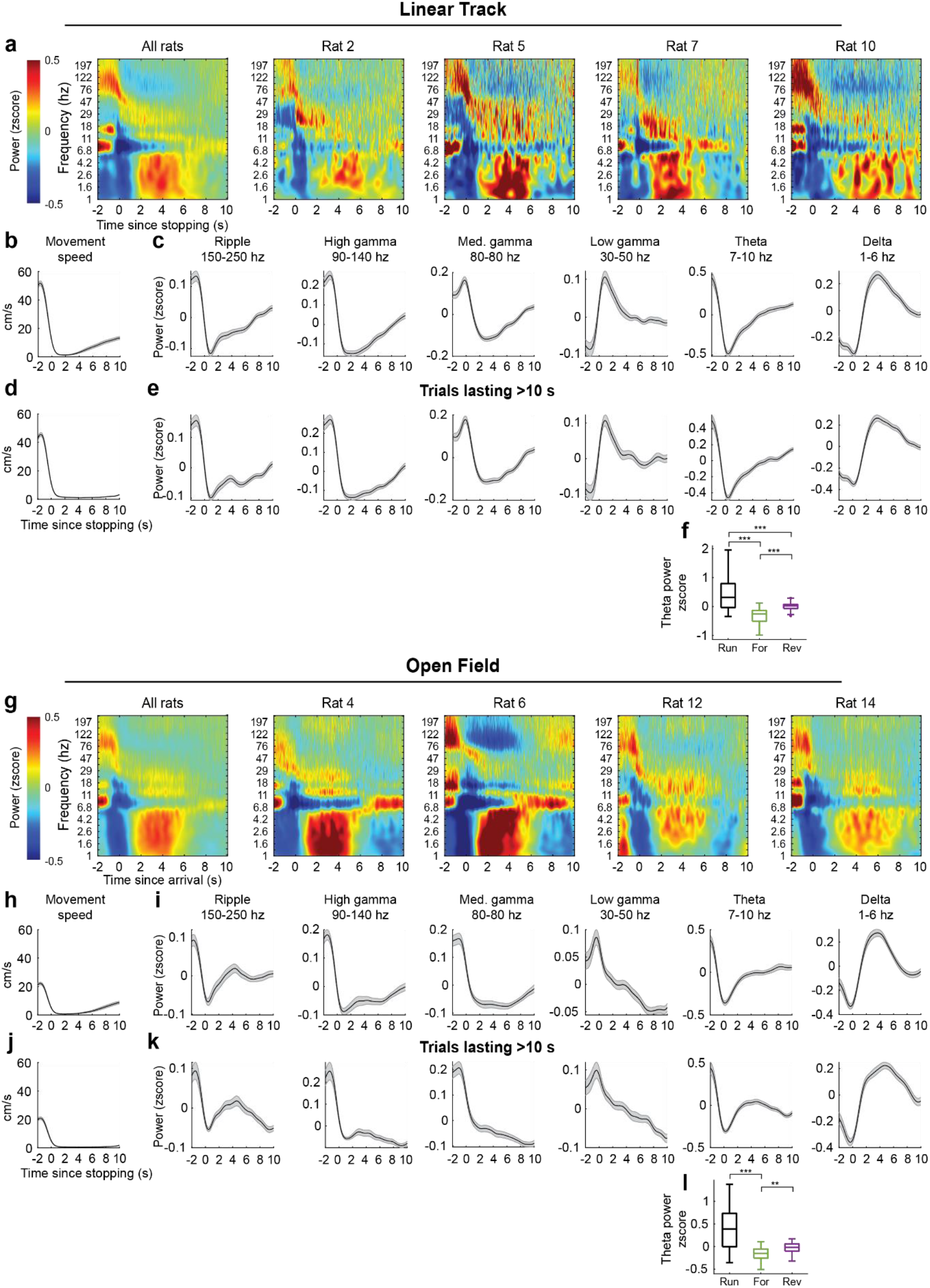
Time-frequency analysis of LFP. **a**, LFP time-frequency spectrograms aligned to stopping (t=0), ranging from -2 (approach to stopping) to +10 seconds. Rats spent variable times drinking, thus later times may include movement (see panels **d-e** for restricted analysis of stops lasting at least 10 s). Left plot shows the average of all linear track sessions (*n=*57 sessions from 10 animals; 1 rat was excluded from LFP analysis due to the presence of large noise artifacts in the LFP). Right plots show session-averaged spectrograms from representative rats. **b**, Movement speed aligned to reward onset (mean±s.e.m. across 57 sessions). **c**, Mean power within different LFP frequency bands (mean±s.e.m. across 57 sessions). For each session the average z-scored power within the frequency range of interest was smoothed with a Gaussian kernel (SD 0.5s). **d**, As in **b**, but for the subset of trials in which the rat drank for at least 10 seconds (mean±s.e.m. across 57 sessions). **e**, As in **c**, but for the subset of trials in which the rat drank for at least 10 seconds (mean±s.e.m. across 57 sessions). **f**, Z-scored theta power during run, the forward window (0-3 s), or reverse window (3-10 s). Box shows the interquartile range, line indicates the median, whiskers indicate the minimum and maximum excluding outliers. The run window was defined as -2 to -1 s before stopping, to capture periods of consistently high velocity. A Friedman test revealed significant differences in theta power between the epochs (*X*^2^(2) = 880.9, *P=*5.3e-192, *n=*57 sessions). Theta power during the forward window was reduced compared to both run and the reverse windows (forward window v run, *P<*0.0001; forward window v reverse window, *P<*0.0001; posthoc Tukey’s honest significant difference tests). **g-i**, As in (**a-c**), but for trials of the open-field navigation task (*n=*47 sessions from 6 rats). **j-k**, As in (**d-e**), but for trials of the open-field navigation task (*n=*47 sessions from 6 rats). **l**, As in (**f**), but for trials of the open-field navigation task (*n=*47 sessions from 6 rats). A Friedman test revealed differences in theta power between epochs (*X*^2^(2) = 29.9, *P=*3.2e-7). Theta power during the forward window was reduced compared to both run and the reverse windows (forward window v run, *P<*0.0001; forward window v reverse window, *P=*0.0024; posthoc Tukey’s honest significant difference tests).

**Fig. S2.**
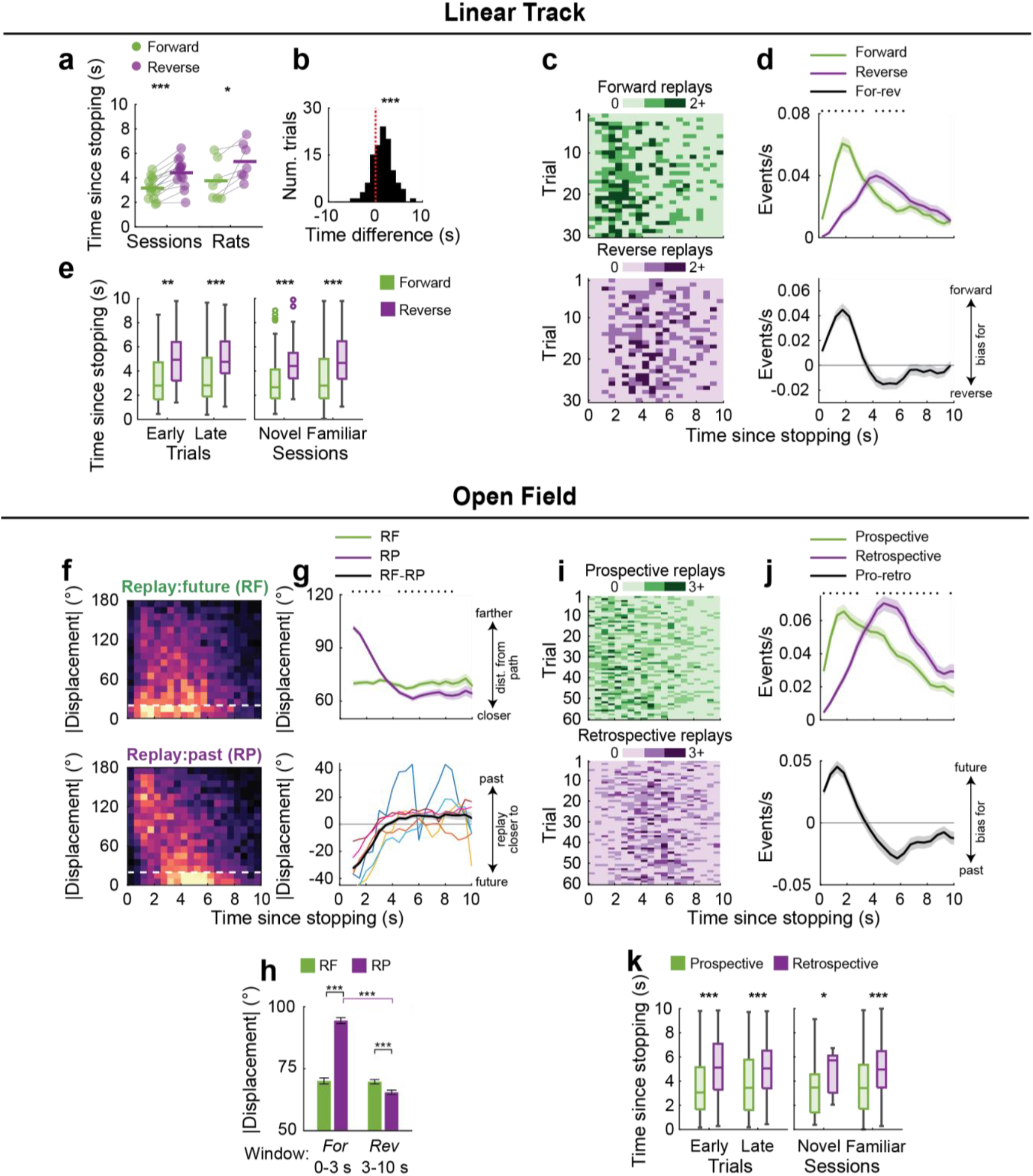
Forward/prospective replays precede reverse/retrospective replays (replays overlapping with sharp-wave ripples). **a-e**, Timing of forward and reverse replays on the linear track, using sharp-wave ripples as candidate replay events. Overall, forward replays preceded reverse replays (*n*=384 forward, 295 reverse replays; median [IQR] time since stopping, forward: 2.7 [1.7-4.5] s; reverse: 4.5 [3.4-6.2] s; *Z=*-9.5, *P*=2.5e-21, Wilcoxon Rank Sum test [WRS]). **a**, Median time of forward or reverse replays (sessions: *n=*14, *P*=2.4e-4, subjects: *n=*7, *P*=0.016, Wilcoxon Signed Rank test [WSR]; includes sessions or subjects with at least 5 forward and 5 reverse replays detected). **b**, Time differences between the first forward and reverse replays within individual stopping periods (*n=*91, median [IQR] difference: 1.7 [0.3-2.8] s; *Z*=5.9, *P*=4.5e-9, WSR). **c**, Replay counts, summed across sessions. **d**, Forward and reverse replay rates (mean±s.e.m., *n=*maximum of 1625 stopping periods; this number decreases over time due to variability in the time spent drinking). **e**, *Left*: Time of forward and reverse replays on early trials (1-10) or late trials (30-40) (early: *n=*42 forward, 43 reverse replays, *Z*=-3.2, *P=*0.0012; late: *n=*84 forward, 60 reverse replays, *Z*=-4.6, *P=*3.8e-06; WRS). *Right:* Time of forward and reverse replays in novel (first exposure) or familiar environments (novel: *n=*132 forward, 87 reverse replays, *Z*=-6.1, *P=*9.6e-10; familiar: *n=*252 forward, 208 reverse replays, *Z*=-7.3, *P=*3.6e-13; WRS). **f-k**, Timing properties of open field prospective and retrospective replays overlapping with a sharp-wave ripple event. **f**, Replay counts (max=40 replays) as a function of angular displacement from future (*top*, RF) or past (*bottom*, RP) paths over time. Dashed line: 20 deg. **g**, Mean±s.e.m. angular displacement from the future or past path (*top*) and difference in angular displacement (*bottom*) (*n=*5,046 replays). Colors: individual rats. Black: rat average. **h**, Angular displacements within the forward (*n*=1,796 replays) or reverse windows (*n*=3250 replays) (interaction *P*=1.0e-4, shuffle; For. window, RF v RP: *P*=4.5e-193; Rev window, RF v RP: *P*=1.2e-10; RF, For v Rev window: *P*=0.086; RP, For v Rev window: *P*=2.0e-315; WRS). **i**, Prospective and retrospective replay counts, summed across sessions. **j**, Rates of prospective and retrospective replays (mean±s.e.m., *n=*maximum of 3095 stopping periods; this number decreases over time due to variability in the time spent drinking). **k**, *Left*: Time of prospective and retrospective replays on early or late trials (early: *n=*79 prospective, 75 retrospective replays, *Z=*-4.0, *P*=7.1e-5; late: *n=*127 prospective, 93 retrospective replays, *Z=*-4.2, *P=*2.7e-5). *Right*: time of replays in novel or familiar environments (novel: *n=*23 prospective, 12 retrospective replays, *Z=*-2.1, *P=*0.033; familiar: *n=*767 prospective, 752 retrospective replays, *Z=*-11.0, *P=*4.5e-28; WRS). Box plots show the median and interquartile range, with whiskers indicating the minimum and maximum. Asterisks in panels **d, g, j** indicate *P*<0.05, WSR tests. *P<*0.05* *P<*0.01** *P<*0.0001***

**Fig. S3.**
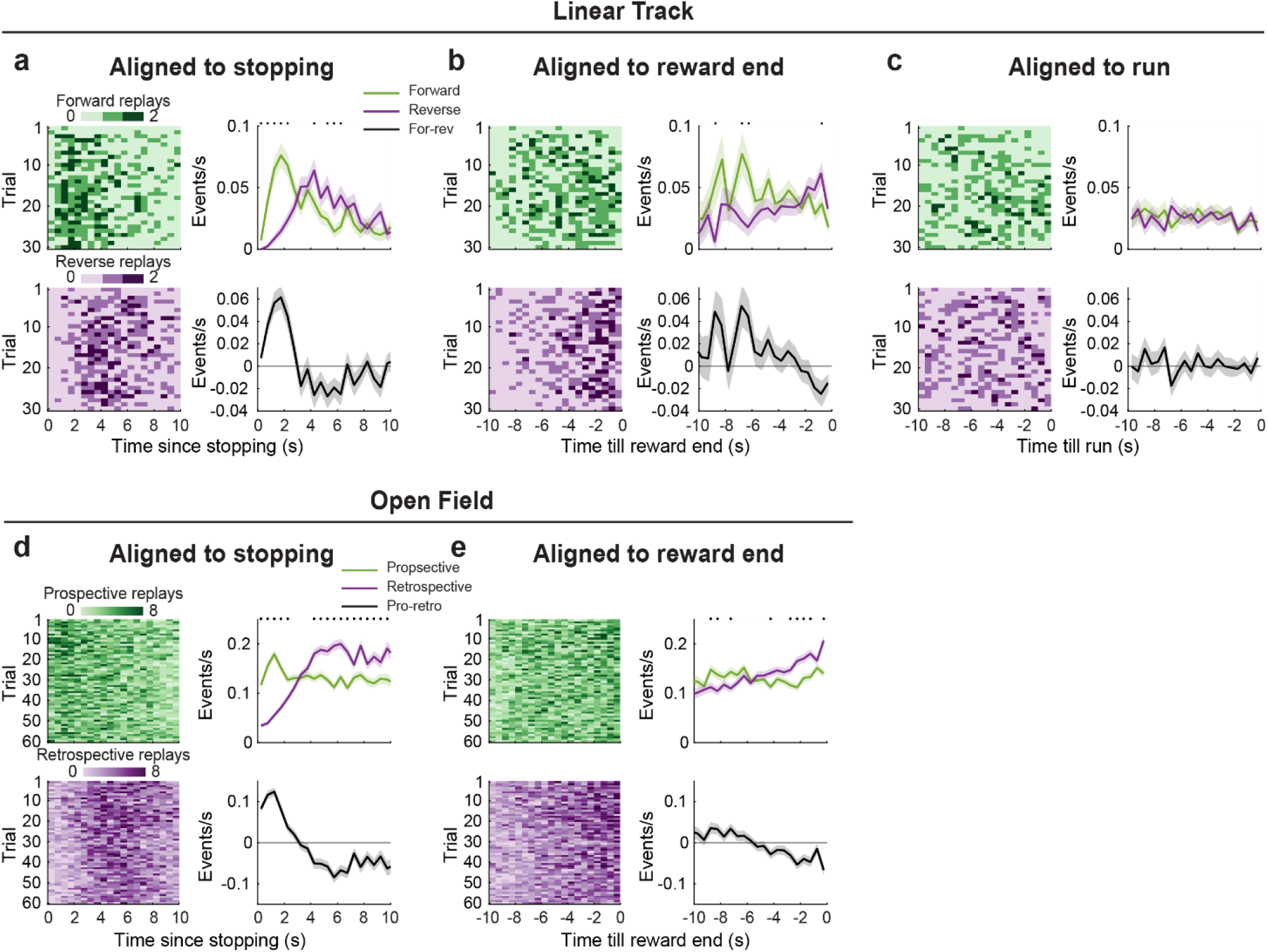
Alignment of replays relative to different behavioral transitions. Fig. 2 shows forward/reverse (linear track) and prospective/retrospective (open field) replay rates aligned to stopping/the start of reward consumption. Here, we additionally show rates aligned to the end of reward consumption (**b, e**) or to run onset (**c**). Panels **a** and **d** show replays aligned to stopping/reward start (as in Main Figures 2–4) for comparison. Note that here we present unsmoothed replay rates to avoid edge artifacts at the boundaries (0.5 second, non-overlapping bins). Overall, we observed more consistent alignment of replays relative to stopping compared to other behaviors. Unlike^1^, we did not find a tendency of forward replay to increase prior to departure. **a**, *Left*: Forward and reverse replay counts, summed across sessions. Color bar indicates number of replays. *Right top*: Forward and reverse replay rates aligned to stopping (mean±s.e.m., *n=*maximum of 1625 stopping periods; this number varies due to variability in the time spent drinking*). Right bottom*: Difference in replay rate (forward-reverse). **b**, As in **a**, but aligned to the end of reward consumption. **c**, As in **a**, but aligned to run onset. **d**, *Left*: Prospective and retrospective replay occurrences in the first 10 seconds after stopping/reward start, over the first 60 open field trials, summed across sessions. Color bar indicates number of replays. *Right top*: Prospective and retrospective replay rates aligned to stopping (mean±s.e.m., *n=*maximum of 4269 stopping periods; this number varies due to variability in the time spent drinking). *Right bottom*: Difference in replay rate (prospective-retrospective). **e**, As in **d**, but aligned to the end of reward consumption. Asterisks indicate *P*<0.05, WRS.

**Fig. S4.**
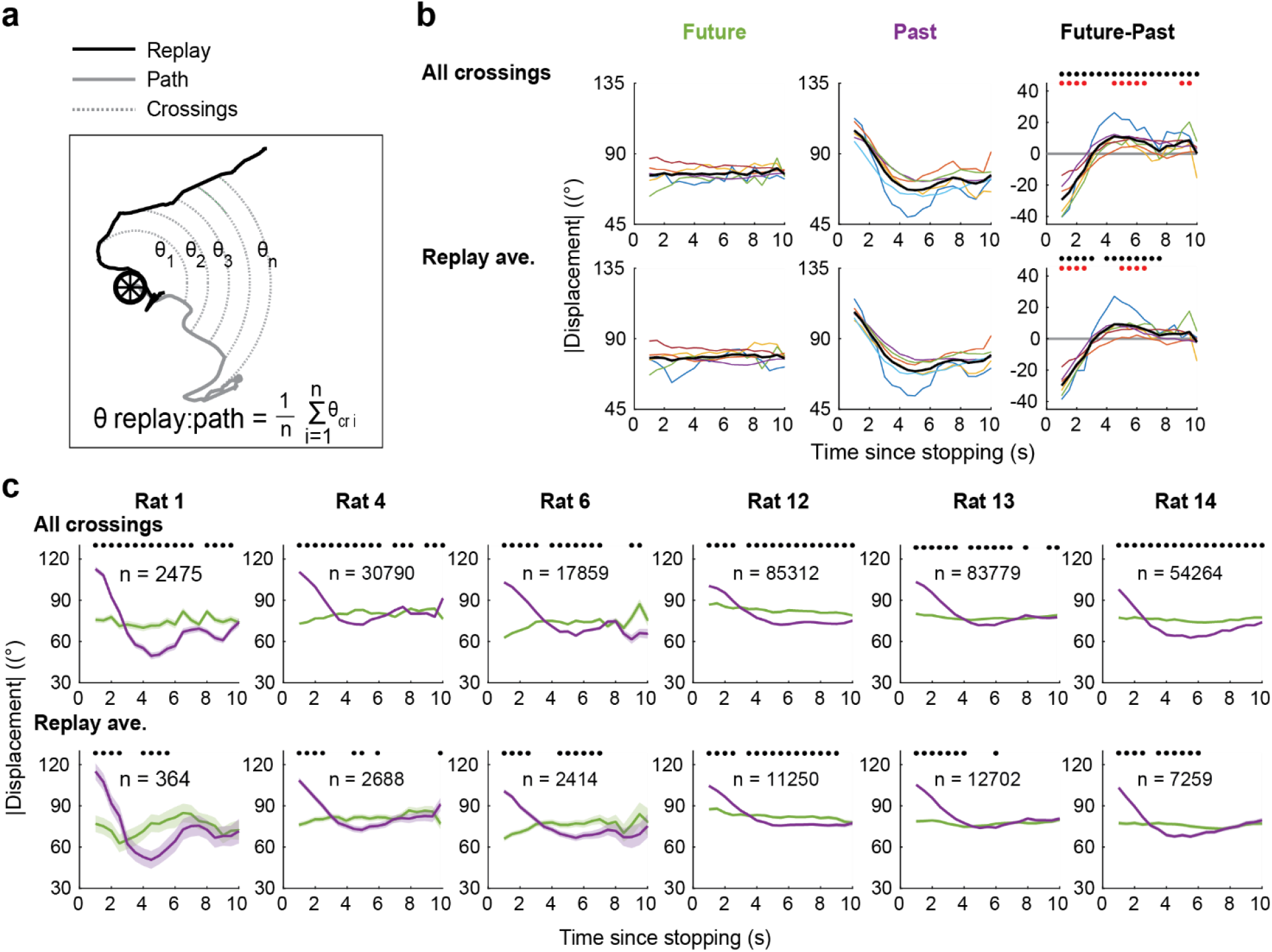
Temporal organization of replay in the open field, individual animals. **a**, Alignment between replayed trajectories and a physically traversed path (future or past) was quantified as the absolute angular displacement between the decoded replay and the physically traversed path at a series of concentric circles with increasing radii, centered at the rat’s current location. In the main text we averaged all the absolute angular displacements to obtain a single value per replay. This allowed for categorization of individual replay events into discrete categories (i.e., ‘prospective’ or ‘retrospective’). Here, we additionally quantify how individual absolute angular displacements (‘crossings’) vary across time, as previously^2^. **b**, Absolute angular displacement of all crossings (*top*) or averaged per replay (*bottom*) relative to the future (*left*) or past (*middle*) path. The difference in angular displacement (future-past) is shown at right. Colored lines represent the averages for individual animals (*n=*6). Black lines represent the average of individual animal traces. Red asterisks and black asterisks at right indicate time bins in which the angular displacements relative to the past or future paths differ significantly (red: WRS tests with *n=*6 animals; black WRS tests with *n=*total number of crossings or replays per time bin). **c**, As in **b**, but plotting the mean±s.e.m. angular displacements per animal. Asterisks indicate time bins in which the angular displacements relative to the future and past paths differed significantly (WRS tests, *P<*0.05).

**Fig. S5.**
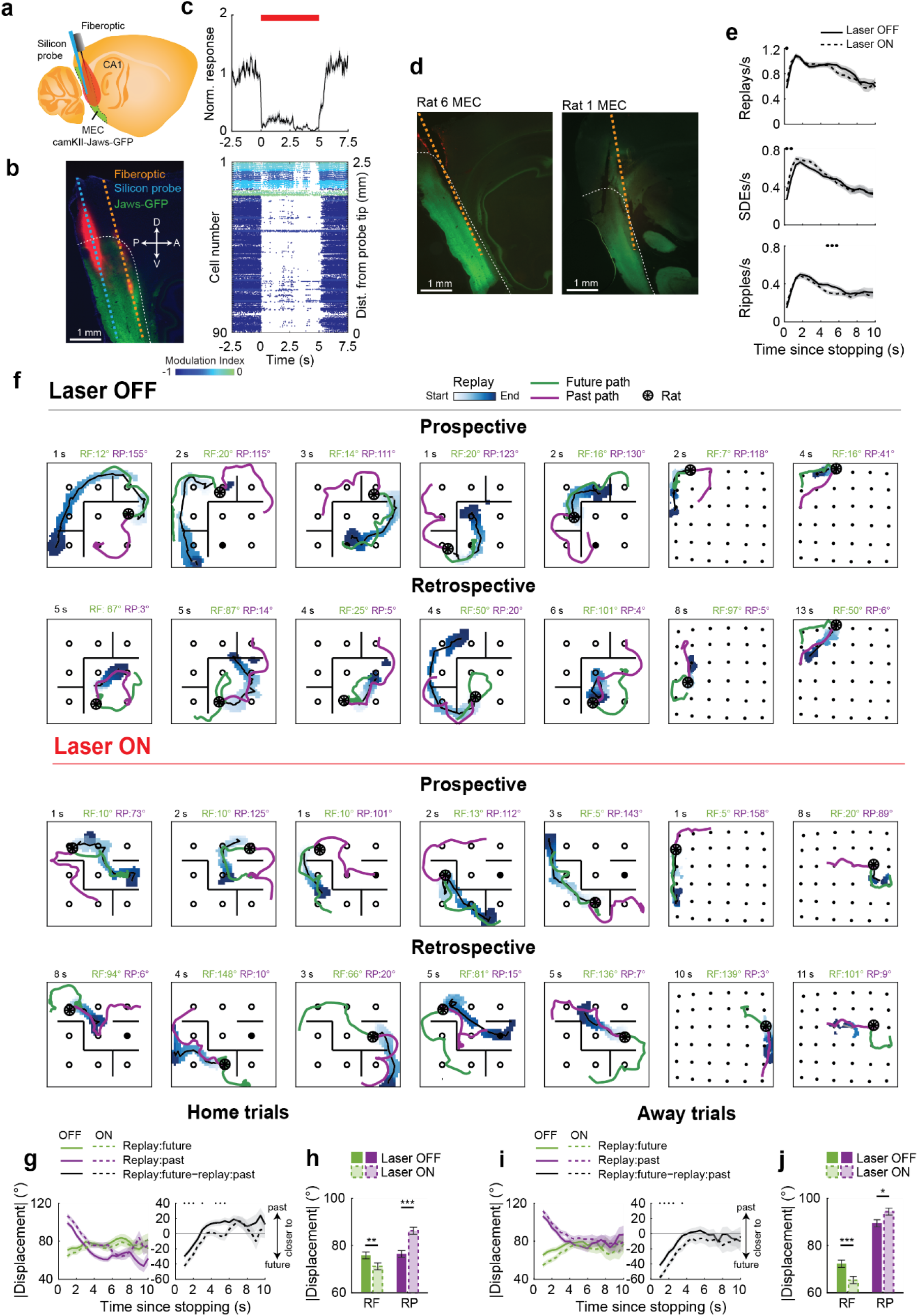
Validation of MEC inhibition and additional analysis of MEC inhibition in the open field. **a-c**, Validation of MEC-inhibition. Similar results were obtained from recordings in 4 hemispheres of 3 rats. **a**, Schematic of the experimental paradigm validating MEC-inhibition in anesthetized animals (Methods). **b**, Histology from an example recording. Image shows expression of Jaws-GFP within MEC, and placement of the tapered fiberoptic (orange) and silicon probe (blue) along the dorsal-ventral MEC axis. A-anterior, P-posterior, D-dorsal, V-ventral. **c**, *Bottom:* Raster plot showing spiking of 90 MEC neurons surrounding 5-second laser pulses (red bar). Color indicates the level of spike modulation (Methods). Spikes from 16 trials are stacked vertically. Cells are organized by their distance from the probe tip, where 0 is positioned most ventrally within MEC. *Top:* Mean±s.e.m. response of the cells below (*n=*90), normalized to pre-laser firing rates. In this recording the firing rate of 81/90 cells was significantly reduced by light delivery (*P<*0.05 in WRS test of firing rates with laser ON versus OFF). **d**, As in Figure 4b. Histology from two additional rats showing Jaws-GFP expression and fiberoptic placement in MEC. **e**, Rate of all replays, spike density events (SDEs) or sharp-wave ripples detected over the first 10s of reward consumption in laser OFF or laser ON sessions (mean±s.e.m., *n=*maximum of 490 laser OFF stopping periods, 498 laser ON stopping periods; this number decreases over time due to variability in the time spent drinking). **f**, Example replays from animals expressing Jaws-GFP in bilateral MEC with laser OFF or ON. The time of the replay since stopping is at top left (black). The angular displacement between the replay and the future (RF) and past (RP) paths are top right. **g-h**, As in Fig. 4g-h, but limited to Home trials. **g**, Angular displacements from past or future paths (*left*) and difference in angular displacement (*right*) on Home trials (*n*=1427, OFF, 1440 ON replays). **h**, Angular displacements within the first 10 seconds of stopping of Home trials (interaction: *P=*1.0e-4, shuffle; OFF v ON, RF: *P=*0.0093, RP: *P=*1.3e-6, WRS tests). **i-j**, As in Fig. 4g-h, but limited to Away trials. **i**, Angular displacements from past or future paths (left) and difference in angular displacement (right) on Away trials (*n*=1351, OFF, 1321 ON replays). **j**, Angular displacements within the first 10 seconds of stopping on Away trials (interaction: *P=*3.0e-3, shuffle; OFF v ON, RF: *P=*5.7e-4, RP: *P=*0.026, WRS tests). Asterisks in **g** and **i** indicate time bins for which *P*<0.05, WRS tests. \**P*<0.05, \*\**P*<0.01, \*\*\**P*<0.001.

**Fig. S6.**
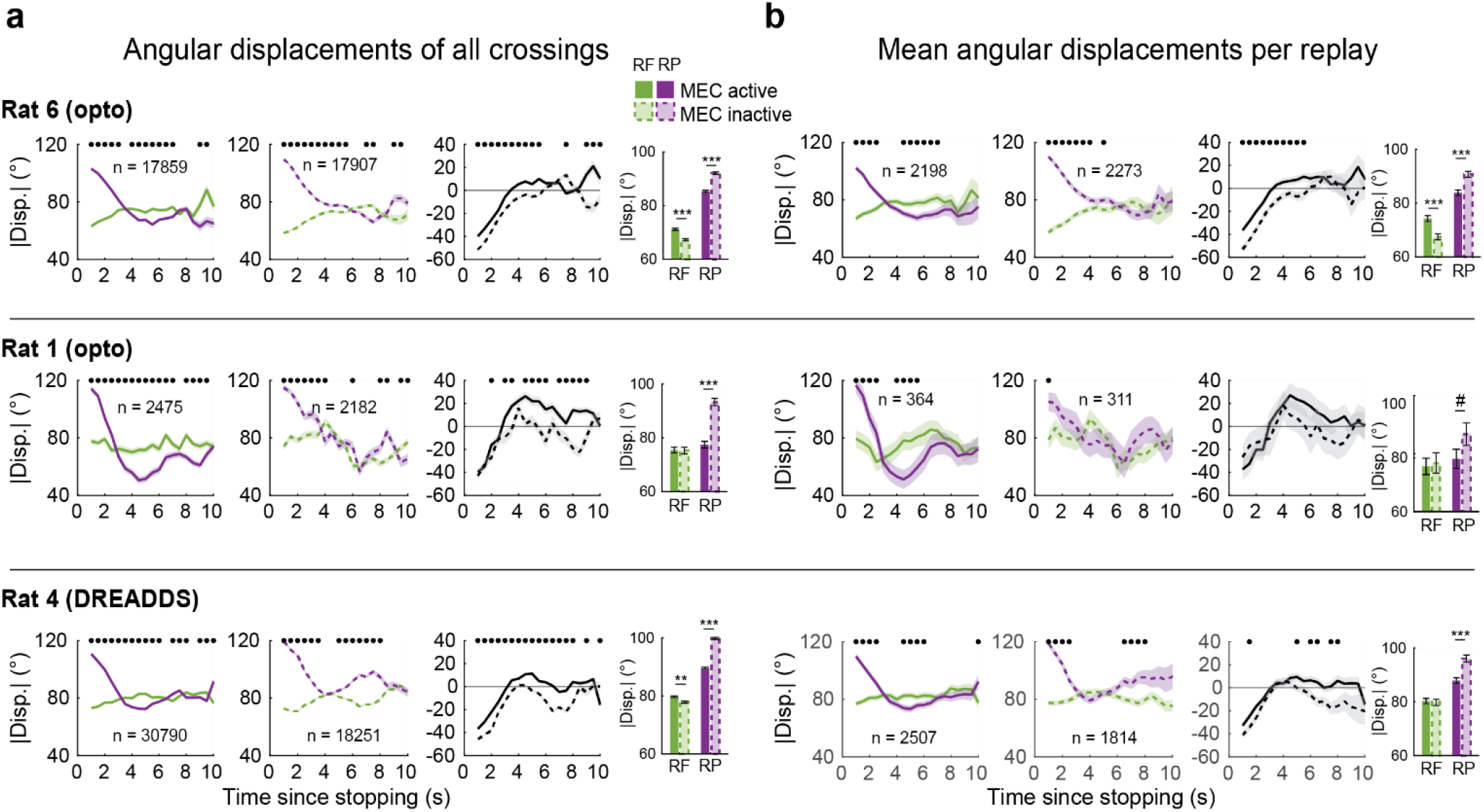
MEC inhibition in the open field, individual subjects. **a**, Angular displacement between replay trajectories and past (RP) or future paths (RF) at each ‘crossing’ point^2^ (see Fig. S4 for schematic of crossings). *Left*: Angular displacements with MEC active (laser OFF sessions for rats 1 & 6, who expressed Jaws-GFP in bilateral MEC and saline sessions for rat 4, who expressed hSyn-hM4D(Gi)-MCherry in bilateral MEC). N denotes the total number of crossings within the first 10 s of stopping. Middle: angular displacements with MEC inactive. *Right*: Difference in angular displacement (RF-RP), with MEC active (solid) versus inactive (dashed). *Far right*: Mean±s.e.m. angular displacements relative to the future or past path, collapsed across the first 10 s of stopping (Rat 6: interaction *P*=6.4e-4, shuffle, RF ON v OFF, *P*=6.4e-14, RP ON v OFF: *P*=3.5e-13, WRS; Rat 1: interaction *P*=6.4e-4, shuffle, RF ON v OFF, *P*=0.41; RP ON v OFF: *P*=3.3e-18, WRS; Rat 4: interaction *P*=6.4e-4, shuffle, RF saline v CNO, *P*=0.004; RP ON v OFF: *P*=2.1e-97, WRS). **b**, As in **a**, but averaging the angular displacements within each replay event. N denotes the total number of replays within the first 10 s of stopping (Rat 6: interaction *P*=6.4e-4, shuffle, RF ON v OFF, *P*=6.7e-6, RP ON v OFF: *P*=3.0e-5,WRS; Rat 1: interaction *P*=0.32, shuffle, RF ON v OFF, *P*=0.86; RP ON v OFF: *P*=0.085, WRS; Rat 4: interaction *P*=3.0e-3, shuffle, RF saline v CNO, *P*=0.67; RP ON v OFF: *P*=8.8e-7, WRS). Asterisks on time course data indicate time bins for which *P*<0.05, WRS. #*P<*0.1, \*\**P*<0.01, \*\*\**P<*0.001

**Fig. S7.**
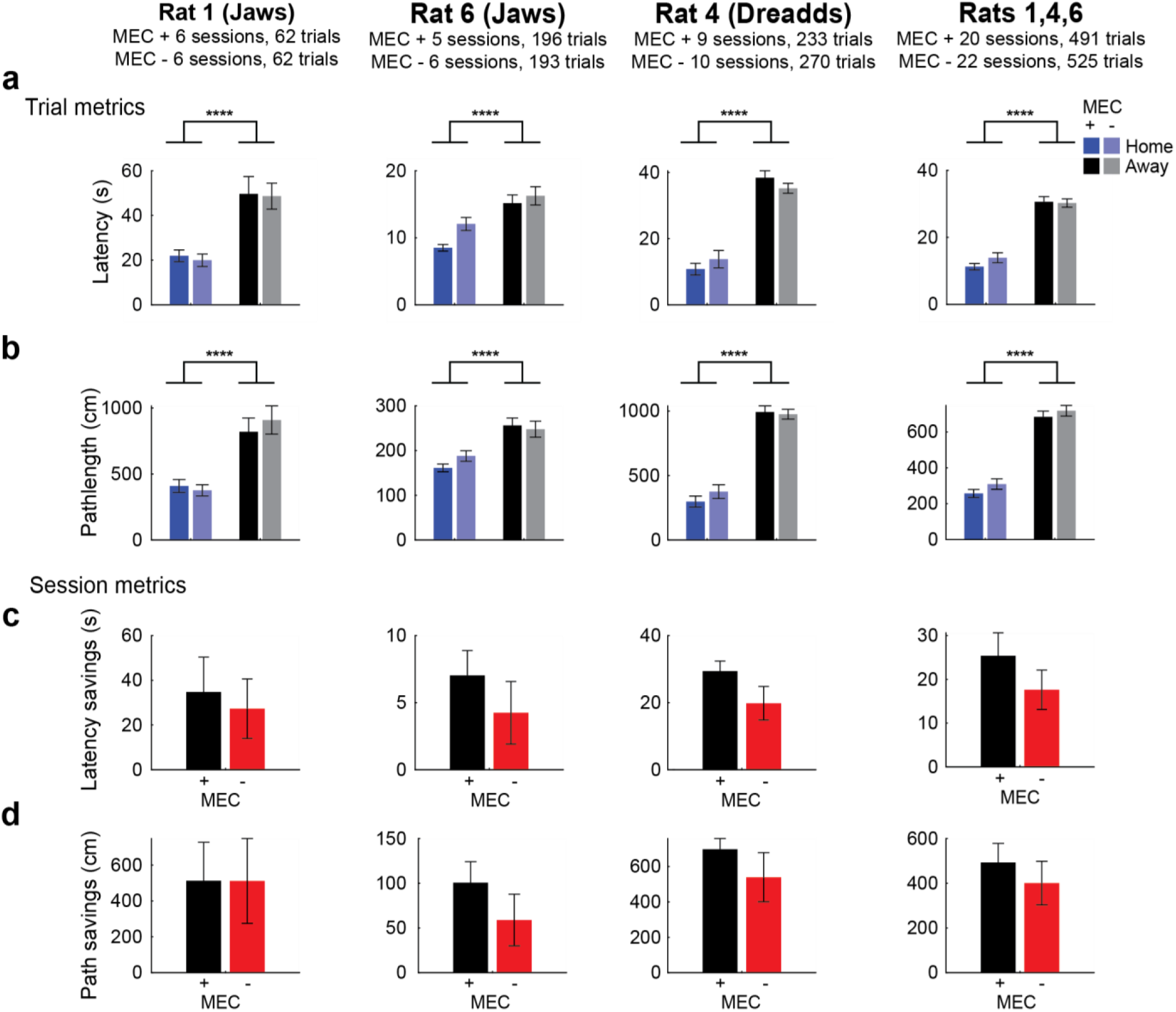
MEC inhibition did not impact performance on the goal-directed navigation task. The open field dataset includes 3 newly recorded rats (with or without MEC inactivation), and 3 rats from a previous publication^3^. Behavioral analysis of the previously published animals can be found in^3^. Here, we show that the 3 newly recorded rats learned the location of the goal on the goal-directed navigation task, and that MEC inactivation did not impair performance. **a**, MEC inactivation did not impact trial latencies. For each rat separately, or all rats combined, we performed a 2-Way ANOVA with MEC activity state (active [+] or inactive [-]) and trial type (Home or Random). For each rat, this analysis revealed a main effect of trial type, indicating that all rats learned the task and remembered the location of the Home well (Rat 1: *F*(1,242)=21.26, *P<*0.0001; Rat 4: *F*(1,991)=142.3, *P<*0.0001; Rat 6: *F*(1,767)=19.29, *P<*0.0001; Rats combined: *F*(1,2008)=106.8, *P<*0.0001). For no rats was there a significant main effect of MEC activity state (Rat 1: *F*(1,242)=0.09, *P*=0.77; Rat 4: *F*(1,991)=0, *P*=0.94; Rat 6: *F*(1,767)=1.1, *P*=0.28; combined: *F*(1,2008)=0.03, *P*=0.87), or a significant interaction between MEC activity state and trial type (Rat 1: *F*(1,242)=0.04 *P*=0.83; Rat 4: *F*(1,991)=1.7, *P*=0.19; Rat 6: *F*(1,767)=3.4, *P*=0.066; combined: *F*(1,2008)=1.9, *P*=0.16). **b**, Pathlength to the home well was reduced compared to pathlength to random wells. MEC inactivation did not impact trial pathlengths. For each rat separately, or all rats combined, we performed a 2-Way ANOVA with MEC activity state (active [+] or inactive [-]) and trial type (Home or Random). For each rat, this analysis revealed a main effect of trial type, indicating that all rats learned the task and remembered the location of the Home well (Rat 1: *F*(1,242)=40.7, *P<*0.0001; Rat 4: *F*(1,991)=198.2, *P<*0.0001; Rat 6: *F*(1,767)=36.2, *P<*0.0001; combined: *F*(1,2008)=226.8, P<0.00001). For no rats was there a significant main effect of MEC activity state (Rat 1: *F*(1,242)=0.02, *P*=0.88, Rat 4: *F*(1,991)=0.6, *P*=0.46, Rat 6: *F*(1,767)=3.2, *P*=0.072, combined: *F*(1,2008)=1.7, *P*=0.20), or a significant interaction between MEC activity state and trial type (Rat 1: *F*(1,242)=0.11, *P*=0.74, Rat 4: *F*(1,991)=0.9, *P*=0.34; Rat 6: *F*(1,767)=3.2, *P*=0.072; combined: *F*(1,2008)=0.4, *P*=0.51). **c**, For each session, we computed the latency ‘savings’ between Home and Random trials as (mean latency on Random trials - mean latency on Home trials). For each rat, and for all rats combined, we compared savings from MEC active (MEC +) and MEC inactive (MEC -) sessions. In no rats did we observe a significant effect of MEC inactivation on latency saving (Rat 1: *P=*0.24; Rat 4: *P=*0.28; Rat 6: *P=*0.25; combined: *Z=*0.94, *P=*0.35; WRS tests). **d**, For each session, we computed the pathlength ‘savings’ between Home and Random trials as (mean pathlength on Random trials - mean pathlength on Home trials). For each rat, and for all rats combined, we compared savings from MEC active (MEC+) and MEC inactive (MEC-) sessions. In no rats did we observe a significant effect of MEC inactivation on pathlength savings (Rat 1: *P=*0.94; Rat 4: *P=*0.84; Rat 6: *P=*0.66; combined: *Z=*0.29, *P=*0.77; WRS tests). \*\*\*\**P<*0.0001

**Fig. S8.**
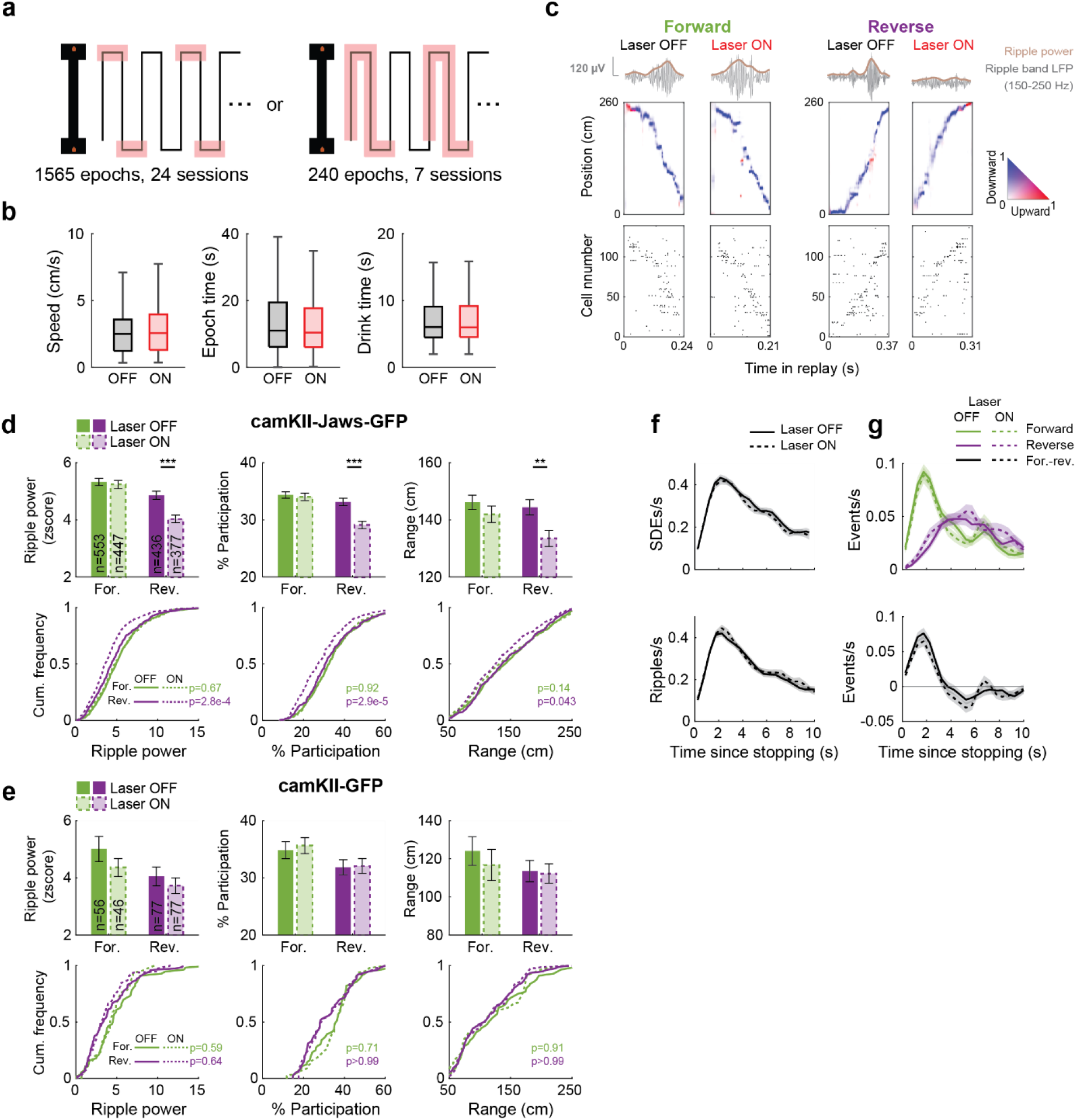
MEC inhibition on the linear track selectively impacted reverse replays. **a**, Experimental paradigm for inhibiting MEC on the linear track. Laser light was delivered to rats expressing camKII-Jaws-GFP (*n=*4) on interleaving laps at the reward platforms (left). In some sessions light was also delivered on the track during run (right). **b**, Rats’ behavior was not significantly impacted by MEC inactivation. N=932 laser OFF and 873 laser ON trials. *Left*: Rat speed on the reward platforms (*Z=*-1.1, *P=*0.25). *Middle*: Time spent on the reward platform (*Z=*0.99, *P=*0.32). *Right*: Time spent drinking reward (*Z=*0.042, *P=*0.98). WRS tests. Box plots indicate the median, interquartile range, and the minimum and maximum excluding outliers, which were omitted for visualization but included in statistical analyses. **c**, Examples of forward and reverse replays from a rat expressing camKII-Jaws-GFP in bilateral MEC, with laser OFF or ON. *Top*: Ripple band-pass filtered LFP (150-250 Hz) and smoothed ripple power during the example events. *Middle*: Posterior probability over position and running direction. *Bottom*: Raster plots of cell spiking during each event. **d**, MEC inactivation reduced the ripple power, cell participation, and spatial coverage of reverse, but not forward, replays. N is reported on bar graphs. *Top panels*: mean±s.e.m. For each property we performed WRS tests between laser OFF and laser ON replays. We tested the significance of the interaction between laser state and replay direction using a shuffle procedure. The observed difference (forward laser OFF-laser ON)-(reverse laser OFF-laser ON) was compared to a distribution of differences obtained by shuffling laser state labels (5000 shuffles, with laser state permuted separately for forward and reverse replays). Ripple power, forward: *Z=*0.58, *P=*0.56; reverse: *Z=*4.2, *P=*2.4e-5, interaction: *P=*0.0076. % participation, forward: *Z=*0.41, *P=*0.68, reverse: *Z=*4.6, *P=*5.0e-06, interaction: *P=*0.0032. Range, forward: *Z=*1.3, *P=*0.21; reverse: *Z=*2.8, *P=*0.0048, interaction: *P=*0.23. *Bottom panels*: cumulative distribution functions. *P*-values from Kolmogorov–Smirnov tests are shown in green (forward Laser OFF versus ON) and purple (reverse Laser OFF versus ON). Axes have been truncated for visualization, but all data was included for statistical tests. **e**, As in **d**, but for laser-control rats expressing camKII-GFP in bilateral MEC (*n=*2 rats). We did not observe significant changes to either forward or reverse replays with light delivery alone. Ripple power, forward: *Z=*0.68, *P=*0.50; reverse: *Z=*0.11, *P=*0.91, interaction: *P*>0.99. % participation, forward: *Z=*-0.48, *P=*0.63, reverse: *Z=*-0.056, *P=*0.96, interaction: *P=*0.84. Range, forward: *Z=*0.68, *P=*0.50; reverse: *Z=*0.11, *P=*0.91, interaction: *P*>0.99. **f**, MEC inactivation did not impact the rate of occurrence of spike density events (SDE, *top left*) or sharp-wave ripples (SWR, *bottom left*). Plots show mean±s.e.m. rates of the first 10 seconds following reward onset (n=a maximum of 947 laser OFF, 900 laser ON epochs; this number decreases over time due to variability in time spent drinking). Overall rates within each stopping period: SDE, OFF 0.26±0.0074 hz, ON 0.26±0.007 hz, *Z=*0.24, *P=*0.8; SWR, OFF 0.23±0.0067, ON 0.23±0.0071 hz, *Z=*0.18, *P=*0.86; forward replay, OFF 0.041±0.0024 hz, ON 0.039±0.0024 hz, *Z=*0.70, *P=*0.49; reverse replay, OFF 0.025±0.0017 hz, ON 0.029±0.0019 hz, *Z=*-1.3, *P=*0.19. WRS tests. **g**, Although MEC inactivation reduced the quality of reverse replays on the linear track (**d**), we did not detect a significant reduction in their rate of occurrence. Top plots show mean±s.e.m. forward and reverse replay rates (*n=*a maximum of 947 laser OFF, 900 laser ON epochs; this number decreases over time due to variability in time spent drinking). Bottom plot shows rate difference (forward-reverse). Overall rates within each stopping period: forward replay, OFF 0.041±0.0024 hz, ON 0.039±0.0024 hz, *Z=*0.70, *P=*0.49; reverse replay, OFF 0.025±0.0017 hz, ON 0.029±0.0019 hz, *Z=*-1.3, *P=*0.19. WRS tests.

**Fig. S9.**
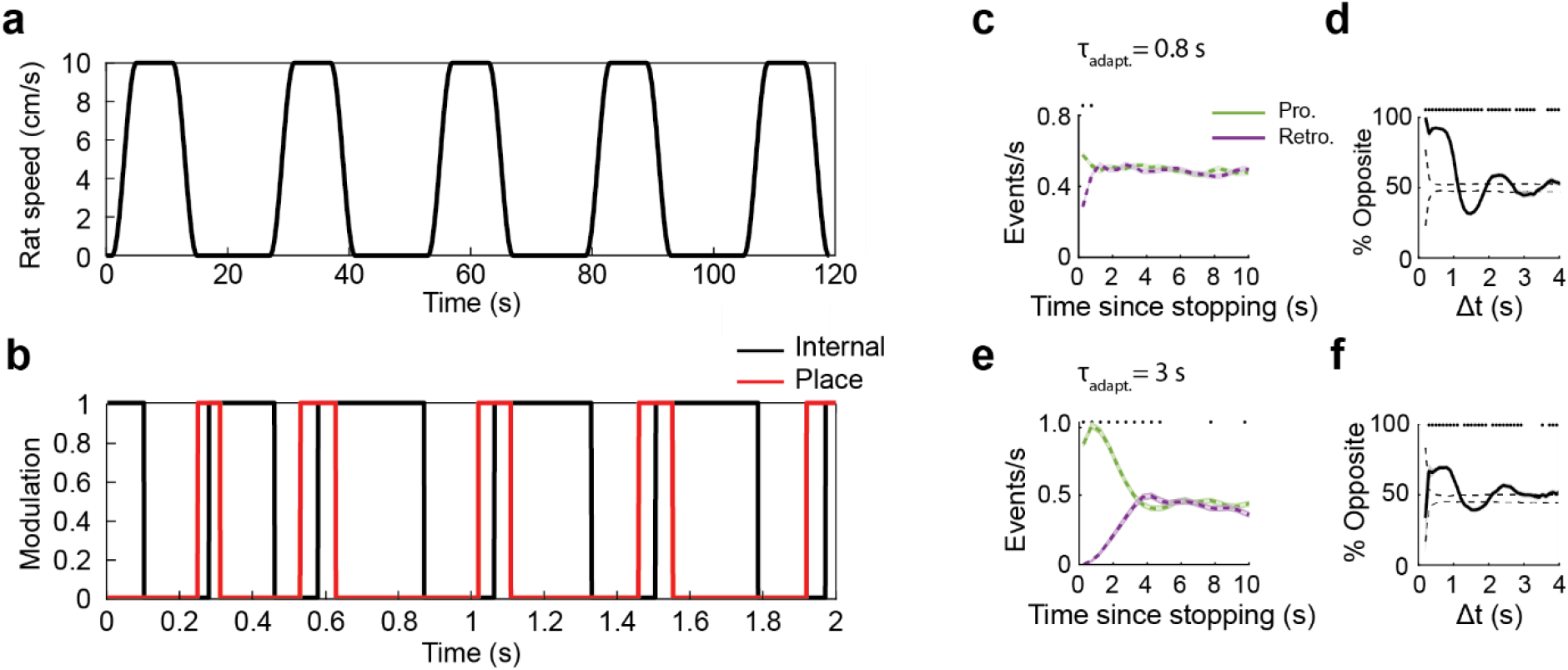
Additional model details. **a**, Simulated rat speed profile across 5 movement bouts. **b**, Theta modulation of ‘internal’ and place inputs across several theta cycles (see Methods). A brief place input was provided to the network at the beginning of each theta cycle, while the internal inputs (recurrent and adaptive) were turned on later in the cycle. **c**, Prospective and retrospective replay rates produced by the adaptation-only model with a shorter time constant (=0.8 s). **d**, Replay self-avoidance, resulting in alternation, produced by model with =0.8 s. **e-f**, As in (**c-d**) but with =3 s (as in the main manuscript). The model produces both past- and self-avoidance with varying time constants of adaptation, but =3 s produces a better match to the observed time course of past-avoidance. Asterisks indicate *P*<0.05, WSR tests.

## Notes

### Competing Interest Statement

The authors have declared no competing interest.

